# Pneumococcal Evasion of Antibiotics via Metabolic Adaptation During Infection

**DOI:** 10.1101/2022.04.14.488313

**Authors:** Tina H. Dao, Haley Echlin, Abigail McKnight, Enolia S. Marr, Julia Junker, Qidong Jia, Randall Hayden, Jason W. Rosch

## Abstract

*Streptococcus pneumoniae* is a major human pathogen of global health concern, causing a range of mild to severe infections, including acute otitis media, pneumonia, sepsis, and meningitis. The rapid emergence of antibiotic resistance among *S. pneumoniae* isolates poses a serious public health problem worldwide. Resistant pneumococcal strains have rendered the mainstay treatment with beta-lactams, fluoroquinolones, and macrolides, ineffective. Antibiotic resistance in *S. pneumoniae* has spread globally via the emergence of *de novo* mutations and horizontal transfer of resistance. Fluoroquinolone resistance in *S. pneumoniae* is an intriguing case because the prevalence of fluoroquinolone resistance does not correlate with increasing usage, as is often the case with other classes of antibiotics. In this study, we demonstrated that deleterious fitness costs constrain the emergence of individual fluoroquinolone resistance mutations in either topoisomerase IV or gyrase A in *S. pneumoniae.* Generation of double point mutations in the target enzymes in topoisomerase IV and gyrase A conferred high-level fluoroquinolone resistance while restoring fitness comparable to the sensitive wild-type. During an *in vivo* model of antibiotic resistance evolution, *S. pneumoniae* was able to circumvent deleterious fitness costs imposed by resistance determinants through development of antibiotic tolerance through metabolic adaptation that reduced the production of reactive oxygen species, an effect that could be recapitulated pharmacologically. The metabolic mutants conferring tolerance resulted in a fitness benefit during infection following antibiotic treatment with fluroquinolones. These data suggest that emergence of fluoroquinolone resistance is tightly constrained in *S. pneumoniae* by host fitness tradeoffs and that mutational pathways involving metabolic networks to enable tolerance phenotypes may be an important contributor to the evasion of antibiotic mediated killing.

## INTRODUCTION

*S. pneumoniae* is a major cause of morbidity and mortality particularly in young children and elderly populations ^1^. Annually, about half a million deaths among children younger than the age of 5, the immunocompromised, and the elderly result from pneumococcal disease worldwide ^2^. Although *S. pneumoniae* usually colonizes the upper respiratory tract asymptomatically, it can become pathogenic especially when host defenses are weakened or compromised, causing mild to invasive diseases, such as sinusitis, acute otitis media, pneumonia, bacteremia, and meningitis ^1, 3^. There has been a worldwide increase in the number of cases of *S. pneumoniae* resistant to multiple classes of antibiotics, including fluoroquinolones (levofloxacin, ciprofloxacin, and gemifloxacin), folate inhibitors (trimethoprim and sulfamethoxazole), cell-wall synthesis inhibitors (penicillin), and protein synthesis inhibitors (azithromycin, doxycycline, tetracycline, and minocycline) ^4^. According to the CDC, in 2015, 30% of about 30,000 cases of invasive pneumococcal disease resulted from *S. pneumoniae* that was resistant to more than one antibiotic ^5^. In the United States, approximately 20% to 40% of pneumococcal isolates are resistant to macrolides ^6, 7^ and 13% to 30% are resistant to beta- lactams ^6, 8–10^. Worldwide, the pneumococcus is the fourth leading pathogen responsible for global deaths associated with antimicrobial resistance, only surpassed by *Escherichia coli*, *Staphylococcus aureus*, and *Klebsiella pneumoniae* ^11^. Following introduction of the Prevnar-13 conjugate vaccine, there have been reports of rising rates of antibiotic resistant pneumococcal infections in pediatric patients ^12^. These data underscore the importance of understanding the mechanisms and constraints that underlie the emergence of antibiotic resistance in *S. pneumoniae*.

Unlike macrolide and beta-lactam resistance, there have been paradoxical observations in the case of fluoroquinolone resistance. Globally, fluoroquinolones are one of the most frequently prescribed antibiotics ^13^. Despite a high usage rate, only limited cases of highly fluoroquinolone-resistant pneumococcal isolates have been reported ^14, 15^. The prevalence of fluoroquinolone resistance remains low regardless of the frequent usage of fluoroquinolones in Asia, Europe, and Spain, ranging from 0.6% to 9.8% (**Table 1**). The highest prevalence of fluoroquinolone resistance (4-13%) was reported in Croatia and Hong Kong ^16, 17^. Unlike the relatively widespread macrolide and to a lesser extent beta-lactam resistance, fluoroquinolone resistance in *S. pneumoniae* remains very low with only 1 to 2% of pneumococcal isolates in the United States being resistant ^13, 18^. These data suggest there may genetic or fitness related constraints for fluoroquinolone resistance in *S. pneumoniae*.

Ciprofloxacin was the first fluoroquinolone prescribed to treat lower respiratory tract infections ^19^. Currently, ciprofloxacin is not the first-line treatment for pneumococcal infection due to its limited potency against *S. pneumoniae* and potential for rapid emergence of resistance ^16, 20–23^. A newer fluoroquinolone, levofloxacin, which has greater potency against *S. pneumoniae*, is one of the current standard treatments for community-acquired pneumococcal pneumonia ^24^. Acquisition of levofloxacin resistance in *S. pneumoniae* has been reported to occur rapidly during antibiotic therapy, with multiple instances demonstrating the emergence of resistance during treatment of invasive disease ^25^. Fluoroquinolones (such as ciprofloxacin, levofloxacin, and moxifloxacin) inhibit topoisomerase IV (ParC and ParE) and gyraseA (GyrA), which are responsible for unwinding supercoiled DNA, thereby halting DNA replication and resulting in cell death ^26^. Mutations in the on-target genes *parC* and/or *gyrA* confer high level resistance to fluoroquinolone resistance in *S. pneumoniae*. The acquisition of fluoroquinolone resistance can occur via horizontal transfer of mutations via recombination with resistant strains or via *de novo* mutation ^27^. The propensity of the pneumococcus for genetic exchange allows for antibiotic resistance determinants to rapidly spread amongst strains which, in theory, would allow for widespread dissemination of fluoroquinolone resistance. Despite this capacity for rapid spread of resistance, strains harboring mutations conferring fluoroquinolone resistance remain relatively rare.

Often overshadowed by the prominent phenomenon of antibiotic resistance, tolerance and heteroresistance are becoming increasingly recognized as being important mechanisms underlying antibiotic treatment failure and recalcitrant infections ^28, 29^. Tolerance refers to the ability of the bacteria to withstand the lethal actions of bactericidal antibiotics with markedly reduced bactericidal kinetics following antibiotic exposure without significant increases in the minimum inhibitory concentration (MIC) ^30^. While antibiotic resistance is readily detected by traditional clinical MIC assays and thus has been extensively studied and characterized, tolerance is often overlooked and undetected. Currently, there is no gold standard for identifying antibiotic tolerant strains. Although antibiotic kill kinetics (or time-kill curves) have been conducted to detect antibiotic tolerance, such assays are laborious and challenging to undertake as part of routine diagnostics ^31^. Despite these obstacles, antibiotic tolerance has been extensively reported in both Gram-negative and Gram-positive bacteria and serve as a potential avenue for the subsequent development of high-level resistance ^32, 33^. Tolerance pathways include stress response, metabolic regulation, transcriptional regulation, efflux / influx regulation, and genes involved in metabolism, transport, and regulation of gene expression ^34–36^. A key factor in whether antibiotic tolerance and resistance emerge and spread within a population is the fitness of such strains. Antibiotic resistance and tolerance are often accompanied with fitness costs such as decreased replication speed ^37, 38^. While those mutations enable the bacteria to survive under antibiotic pressure, they often render the target enzymes suboptimal in the absence of antibiotics and the resistant/ tolerant strains can be effectively outcompeted by sensitive strains once antibiotic pressure is lifted. It is strains that obtain mutations that engender the capacity to survive antibiotic exposure without conferring corresponding fitness tradeoffs that represent the greatest concern for treatment failure. It is becoming increasingly recognized that addressing the host-pathogen interface must include the understanding of the development of tolerance in the presence of innate immune cells and how the evolutionary pathways to tolerance develop in the host, which may be distinct from mechanisms evolved during *in vitro* selection.

Antibiotic treatment failure, whereby initial antibiotic therapy fails to effectively clear the infection, can result in prolonged infections and poorer clinical outcomes. For bacterial nosocomial pneumonias, initial antibiotic treatment failure has been reported as high as >70% of patients and, even when resistant organisms are not present, the mortality rate for hospitalized patients can be great than 50% ^39^. Similar observations are made with complicated urinary tract infections whereby the 40% mortality rate of a multi-drug resistant organism is only reduced to a 30% rate for a susceptible strain ^40^. Discontinuation of initial antibiotic treatment is often due to perceived clinical failure rather than isolation of a multi-drug resistant pathogen ^41^. Traditional antimicrobial testing mechanisms performed during clinical failure only identify truly resistant pathogens and do not identify tolerant populations of bacteria that manifest only under specific growth conditions, such as biofilm growth, or can only be identified using population analysis profiles ^42, 43^. Although not detected by traditional testing, these subpopulations of bacteria recalcitrant to antibiotic killing are increasingly thought to be important contributors to antibiotic treatment failure ^29, 44–46^.

In our current study, we investigated whether there are barriers to fluroquinolone resistance in *S. pneumoniae* that may explain the continued low levels of resistance that are observed clinically. We found that the first step mutations in either *gyrA* or *parC* imparted a high fitness cost during *in vivo* infection while strains harboring multiple mutations regained the capacity to cause invasive disease. In agreement with the attenuation of the individual mutants, experimentally evolved strains subjected to repeated fluroquinolone treatment failed to evolve resistance. These evolved isolates, however, developed tolerance to fluroquinolone mediated killing. This phenotype was linked to the reduced capacity of the evolved strains to produce oxygen radicals in the form of hydrogen peroxide, which, in turn, limited DNA damage during antibiotic treatment. These data indicate that *S. pneumoniae* can evade antibiotic mediated killing via inactivation of specific metabolic networks to enable tolerance phenotypes.

## METHODS

### Media and growth conditions

For this study, *S. pneumoniae* was either grown in C+Y (a chemically-defined semi-synthetic casein liquid media supplemented with 0.5% yeast extract), ThyB media (Todd Hewitt Broth + 0.2% yeast extract; BD), or on blood agar plates containing tryptic soy agar (EMD Chemicals, New Jersey) and 3% defibrinated sheep blood. Cultures of *S. pneumoniae* were inoculated from overnight streaked blood agar plates and then incubated at 37°C, 5% CO2. All blood agar plates were supplemented with neomycin at 20 µg/mL for *S. pneumoniae*, which is naturally resistant to neomycin. For transformations, the respective fluoroquinolone for selection was added to the blood agar plates at concentrations indicated. Strains used in this study are indicated in Supplementary Table 2.

### MIC determination via E-tests

The wild-type TIGR4, single point mutants S79Y *parC* and S81F *gyrA*, and double mutants S79Y *parC* & S81F *gyrA* and S79Y *parC* & D435N *gyrB* strains were grown on blood agar plates overnight, inoculated into ThyB, and incubated at 37°C, 5% CO2 until reaching mid-logarithmic phase at OD620 ∼0.4. Then, 100 µL of the bacterial culture was spread onto TSA blood agar plates. A levofloxacin, ciprofloxacin, or moxifloxacin E-test strip (bioMerieux) was added onto the center of each plate. Minimum inhibitory concentration MIC (µg/mL) was determined to be the concentration at which the symmetrical inhibition ellipse edge intersects the E-test strips on the plate ^47^.

### Genomic DNA extraction

To extract genomic DNA for transformation and whole genome sequencing, the aqueous/organic extraction method using phenol chloroform was performed. TIGR4 and mutant strains were grown in ThyB to late logarithmic phase at OD620 ∼0.6 and pelleted at 6,000 x g for 10 minutes followed by resuspension in 1 mL genomic lysis buffer containing 50 µL 10% DOC, 50 µl 10% SDS, and 10 µL of 10mg/mL Proteinase K (Sigma). The mixture was incubated for lysis until clear (approximately 5 minutes) at 37°C, mixed with 500 µL phenol: chloroform: isoamyl alcohol (Sigma), transferred to phase-lock tubes (Quantabio), and centrifuged at maximum speed for the separation of aqueous and organic phases. Then 500 µL chloroform: isoamyl alcohol (Sigma) was mixed with the aqueous phase and centrifuged as above. For DNA precipitation, the aqueous phase was transferred to a new tube containing 100% ethanol.

Precipitated DNA was washed with ice cold 70% ethanol, pelleted, dried, and resuspended in nuclease-free water.

### Transformation

Cultures of *S. pneumoniae* were inoculated in C+Y media and incubated at 37°C, 5% CO2 until OD620∼0.07 to 0.1. Then, 5 µg of genomic DNA and 3 µL of 1 mg/mL CSP1, CSP2, or both, were added to 1 mL of the bacterial cultures of D39, TIGR4, or CDC001, respectively. Three different types of DNA were used for transformation experiments: 2 kb PCR fragments encoding select point mutations, genomic DNA of fluoroquinolone resistant *S. pneumoniae*, or genomic DNA of *S. viridans* clinical isolates. All clinical isolates were genotyped at the respective resistance loci by PCR and Sanger sequencing. 3 kb PCR fragments were generated by amplifying 1 kb of the upstream and downstream nucleotides flanking the select point mutations in *gyrA* and/or *parC* from fluoroquinolone-resistant TIGR4 strains, which were evolved during *in vitro* passaging. The genomic DNA from both species was extracted using the aqueous/organic extraction protocol as described previously. After addition of DNA and CSP, the cultures were incubated for 3 hours at 37°C, 5% CO2, followed by plating of 100 µL of the transformation mixture onto 5 blood agar plates containing select antibiotics. Colony-forming units (CFUs/mL) were enumerated at 24 and/or 48 hours of incubation. Transformation efficiency was calculated as the ratio of transformant colonies (CFUs/mL) on select antibiotic plates to the total number of colonies (CFUs/mL) on blood plates without the antibiotic.

### *in vitro* passaging

TIGR4 was diluted from previously frozen glycerol stocks 1:100 in fresh ThyB containing increasing subinhibitory concentrations of either ciprofloxacin or levofloxacin and incubated until reaching late logarithmic phase (OD620∼0.6). Then, bacterial culture was mixed with glycerol at a final concentration of 20% and stocks were stored at -80°C before serving as an inoculum for the next passage. Each subsequent passage contained progressively increasing 2-fold concentration of the respective antibiotic. The passaging continued for 30 days. Bacterial culture of the last passage at the highest concentration of ciprofloxacin or levofloxacin was spread on blood agar plates, and three colonies were selected for identification of mutations that arose at the end of passaging with either levofloxacin or ciprofloxacin via whole genome sequencing.

### Whole genome sequencing

Genomic DNA samples were sent to Hartwell Center for Bioinformatics and Biotechnology whole genome sequencing. Sequence libraries were prepared using Nextera kits and sequenced according on Illumina HiSeq. All sequences reads are available at NCBI under accession # PRJNA407467. The reads were compared to the TIGR4 reference genome (AE005672.3) using breseq for identification of mutations resulting from recombination events and *de novo* point mutations ^48^.

### Genotype inferences from populations and Muller plots

Mutation filtering, allele frequencies, and plotting were done in R v3.5.3. Muller plots were generated using the lolipop package (https://github.com/cdeitrick/lolipop) v0.6 using default parameters. To summarize, these tools predict genotypes and lineages based on shared trajectories of mutations over time and test their probability of nonrandom genetic linkage. Successive evolution of genotypes, or nested linkage, is identified by a hierarchical clustering method. The method also includes customizable filters that eliminate singletons that do not comprise prevalent genotypes. Muller plots were color coded by the presence of putative driver mutations within each genotype. Additional mutations that occurred on the background of putative driver mutations can be viewed in the allele frequency plots also generated by this package.

### Antibiotic kill curves

The wild-type TIGR4, mutants generated in TIGR4, and the experimentally evolved isolates were inoculated in ThyB at a low cell density OD620 ∼0.05. When cells reached early logarithmic phase (OD620=0.2), the culture was split into untreated and treated conditions. For treatment with antibiotics, the bacteria were exposed to progressively increasing concentrations of ciprofloxacin, levofloxacin, or moxifloxacin, ranging from 1x to 8x MIC of the wild-type (4 µg/mL, 1 µg/mL, and 0.12 µg/mL, respectively). For treatment with edaravone, 3-Methyl-1-phenyl-2- pyrazolin-5-one (Edaravone; Millipore 443300) was resuspended in ethanol at 500mM and heated at 65°C for solubilization. Edaravone was added to the culture at a final concentration of 1mM concurrently with the antibiotics. After addition of treatment, the cultures were incubated at 37°C, 5% CO2 for four hours. For anaerobic conditions, when cells reached early logarithmic phase (OD620=0.2), the culture was pelleted for 5 min, 6,000 x g. Keeping the cellular pellet on ice, the media was deoxygenated by incubation with oxyrase (Oxyrase) at 10% culture volume at 37°C, 5% CO2 for 30 minutes. The cellular pellet was then resuspended in the deoxygenated media, followed by treatment with antibiotics. For all conditions, bacterial cell number was determined hourly by serially dilution of the culture and plating on TSA blood agar plates plus 20 µg/mL neomycin plus 3% defibrinated sheep blood. After incubation overnight at 37°C, 5% CO2, colonies were enumerated and CFU/mL was calculated.

### Growth Curves

Strains were grown to mid-logarithmic phase (OD620 = 0.4) in C+Y and frozen at ^-^80°C in glycerol at a final concentration of 20%. Cells were then thawed and back-diluted 1:100 in 200 µL C+Y in a 96-well plate. Growth at 37°C in 5% CO2 was monitored using Biotek Cytation 3 plate reader for 24 hours.

### TUNEL assay

TIGR4 was inoculated into ThyB at a low cell density (OD620∼0.05) from TSA blood agar plates. When cells reached early logarithmic phase (OD620=0.2), the culture was split into four 25 mLs aliquots plus one 5 mL aliquot. The 5 mL aliquot represents the 0 hour treatment. Each 25 ml aliquot was treated with no antibiotic or antibiotic at 2x MIC of the wild-type and either no edaravone or 1 mM edaravone. After treatment, the cultures were incubated at 37°C, 5% CO2 for four hours. Every hour, 5 mLs of culture was transferred from the 25 mL culture to a new conical tube. 100 µL was used to determine bacterial cell number via serially dilution of the culture and plating on TSA blood agar plates. Each 5 mL of culture was centrifuged 6,000 x g, 5 min and the supernatant was removed. To measure the level of DNA fragmentation via TUNEL staining, the cellular pellet was fixed and stained using a TUNEL kit (BD #556381) via adaptation of the manufacturers protocol. At each timepoint, the cellular pellet was fixed in 1 mL of 1% paraformaldehyde (diluted from 16% paraformaldehyde in PBS) for 1 hour on ice. The pellet was washed with 1x PBS three times and resuspended in 70% ethanol, followed by incubation at -20°C overnight. The pellet was stained according to the manufacturers protocol, with a final resuspension of 300 µL. Levels of FITC (535/623) and PI (488/520) were detected by transferring 200 µL of the resuspended pellet into a black-bottom 96 well plate and measuring fluorescence on a BioTek Cytation 3 plate reader. DNA damage was calculated as the level of FITC (apoptotic cells) divided by the level of PI (total DNA). DNA damage over time was calculated by division of DNA damage at timepoint 1 through 4 hours by the DNA damage at timepoint 0. The experiment was repeated in triplicate.

### H_2_O_2_ production measurement

TIGR4 and the evolved isolates were grown in ThyB until mid-logarithmic phase (OD620 ∼0.4), centrifuged, and resuspended in 1x PBS. After incubation in PBS for 30 minutes, the bacteria were pelleted via centrifugation. The supernatant was collected for the Amplex Red Hydrogen Peroxide/Peroxidase assay (ThermoFisher Scientific #A22188) and 50 µL Amplex Red reagent was added to 50 µL of the supernatant in each microplate well. The absorbance value (OD562) was measured using a BioTek Cytation 3 plate reader. Levels of H_2_O_2_ of each strain was calculated and normalized to its amount of total cellular protein, which was determined via a bicinchonicic acid (BCA) assay (Pierce). The Δ*spxB*Δ*lctO* double mutant was included as a negative control as it produces minimal H_2_O_2_ ^49^. Hydrogen peroxide levels were compared with unpaired parametric t test in Prism 7.

### *in vivo* virulence

To determine the virulence of the mutants in terms of fitness costs *in vivo*, 6-week-old female BALB/c mice (Jackson laboratory) were intranasally infected with the wild-type TIGR4 or the mutant strains (10^6^ CFUs in 100 µL). At 24 hours post challenge, 2 µL of nasal samples and 5 µL of blood samples were collected via nasal lavage and tail bleeding, respectively. The nasal and blood samples were then serially diluted and spotted on TSA blood agar plates for enumeration of bacterial burden. Plates were incubated overnight at 37°C, 5% CO2. Mice were closely monitored for disease progression for 10 days and euthanized via CO2 asphyxiation.

Bacterial titers (CFUs/mL) were compared using non-parametric Mann-Whitney t-test and survival data were analyzed with Mantel-Cox log rank tests in Prism 7.

### *in vivo* passaging

To model the evolution of fluoroquinolone resistance *in vivo*, three groups of 7-week-old female BALB/c mice (Jackson laboratory) were intranasally infected with 100 µL containing 10^6^ CFUs of TIGR4. Six hours post-infection, mice were administered a sublethal dose of levofloxacin intraperitoneally. The initial dosage of levofloxacin (25 mg/kg) was utilized as it eliminated 90% to 99% of the bacterial population (data not shown). After 15 passages, the levofloxacin dosage was increased in a stepwise fashion to increase antibiotic selective pressure. For each passage, twelve hours after levofloxacin treatment, bacteria were collected from lungs of infected mice via dissection. The lungs were homogenized and then bacteria in lung homogenates were collected via centrifugation at 300 x g for 5 minutes, plated on TSA blood agar plates and grown at 37°C, 5% CO2, collected from the agar plates, and frozen in aliquots containing glycerol at a final concentration of 20%. Frozen aliquots from each passage served as the inoculum to infect the next group of mice for the subsequent, respective lineage, and the remaining aliquots were used for genomic DNA extraction.

### *in vivo* kill kinetics

To model the impact of antibiotic tolerance *in vivo*, 8-week-old female BALB/c mice (Jackson laboratory) were intraperitoneally infected with the wild-type TIGR4 and the Δ*spxB*Δ*lctO* mutant strain concurrently (5x10^5^ CFUs of each strain in 100 µL). Sixteen hours post-infection, a blood sample was taken via tail bleeding (timepoint 0). Mice were administered a dose of 1x PBS (N=5) or levofloxacin at 25 mg/kg (N=15) intraperitoneally. Blood samples were taken every 2 hours up to 10 hours via tail bleeding. Bacterial burden in the blood was determined by serial dilution of the blood and spotting on duplicate TSA blood plates containing 20 µg/mL neomycin or 20 µg/mL neomycin plus 1 µg/mL erythromycin. Plates were incubated overnight at 37°C, 5% CO2, colonies were enumerated, and CFU/mL was calculated. The competitive index of Δ*spxB*Δ*lctO* over TIGR4 was calculated as the CFU/mL of Δ*spxB*Δ*lctO* (CFU/mL on the plates containing 20 µg/mL neomycin plus 1 µg/mL erythromycin) divided by the CFU/mL of TIGR4 (CFU/mL on the plates containing 20 µg/mL neomycin minus the CFU/mL on the plates containing 20 µg/mL neomycin plus 1 µg/mL erythromycin). Competitive index of timepoints 2 through 10 were compared to the competitive index of timepoint 0 using unpaired parametric t test with Prism 7.

### Ethics statement

All experiments involving animals were performed with prior approval of and in accordance with guidelines of the St. Jude Institutional Animal Care and Use Committee. The St. Jude laboratory animal facilities are fully accredited by the American Association for Accreditation of Laboratory Animal Care. Laboratory animals are maintained in accordance with the applicable portions of the Animal Welfare Act and the St Jude Animal Care and Use Committee.

## RESULTS

### Fluoroquinolone Resistance Development During Experimental Evolution

Although the prevalence of fluoroquinolone resistance remains relatively low compared to other antibiotics, it still occurs clinically (**Supplementary Table 1**). Cases of levofloxacin resistance in *S. pneumoniae* arising during therapy have been reported in multiple instances ^25^. We first attempted to determine whether such clinical observations for the emergence of resistance could be recapitulated in a murine model of infection and to determine what mutations could arise that would permit resistance. To search for mutations associated with fluoroquinolone resistance *in vivo*, we developed a murine challenge model for studying the emergence of fluoroquinolone resistance using *in vivo* experimental evolution (**Figure 1A**). The wild-type TIGR4 underwent 30 passages in three independent lineages in BALB/c mice under increasing levofloxacin antibiotic dosage (**Figure 1B**). Whole genome sequencing analysis of each experimentally evolved population collected from the final passage indicated the absence of on-target mutations in either *gyrA* or *parC*. In concordance with this observation, minimal shifts in MIC were observed (**Supplemental Table 2**), further supporting that resistance development is absent in these lineages despite repeated antibiotic exposure.

**Figure 1.**
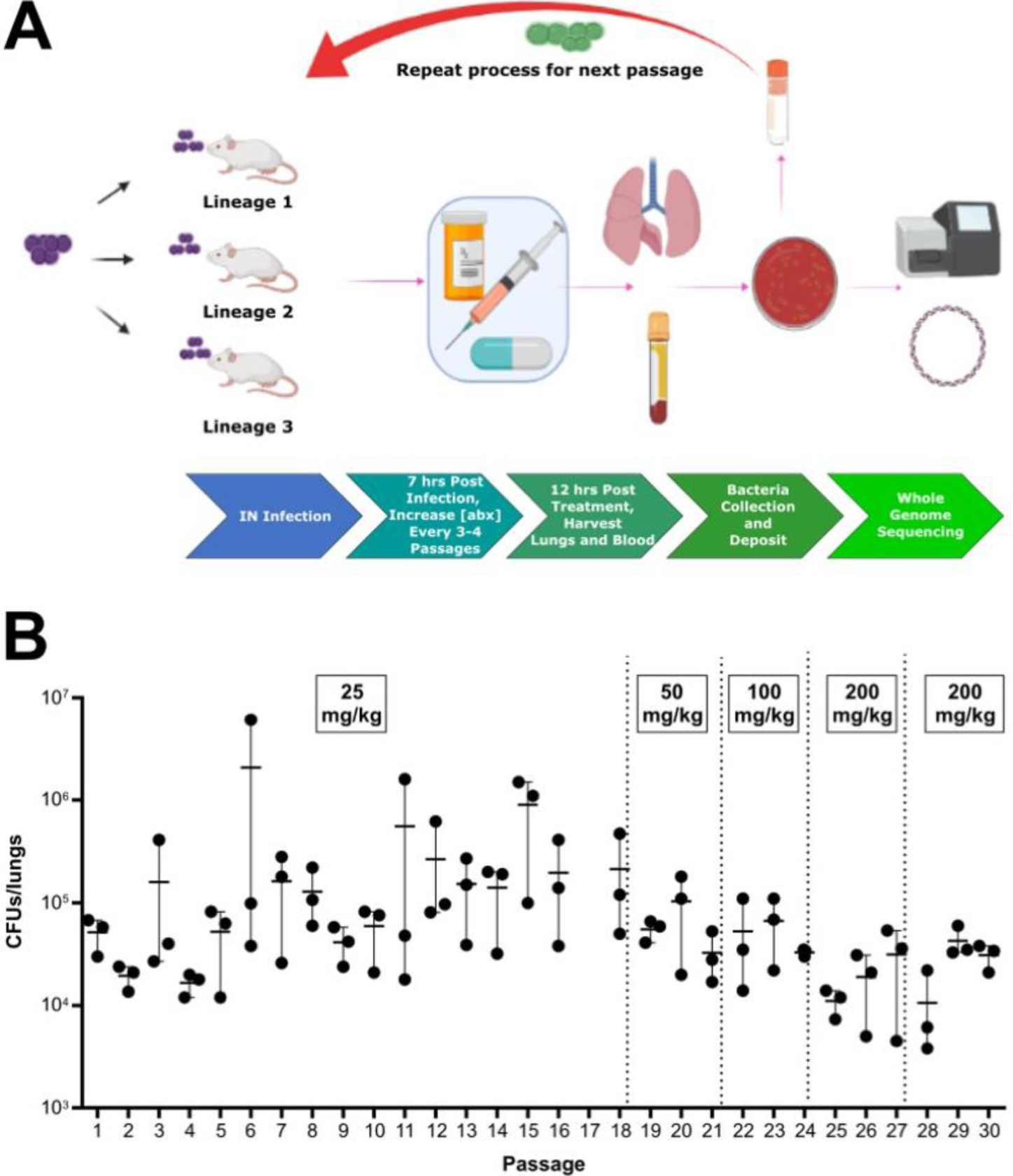
Experimental evolution of *S. pneumoniae* in response to levofloxacin during infection. Schematic diagram of *in vivo* passaging workflow. Mice were intranasally infected with TIGR4 and, after six hours of infection, mice were treated with levofloxacin. Twelve hours after levofloxacin treatment, bacteria were collected from lungs of infected mice and recovered on media plates following overnight incubation. Frozen aliquots from each passage served as the inoculum to infect the subsequent group of mice for each lineage. Created with BioRender.com (A). After each passage under levofloxacin pressure, bacterial burden in the lungs was enumerated. Dosage of levofloxacin ranged from 25 mg/kg to 200 mg/kg as indicated. Each datapoint represents an individual mouse (B).

As the *in vivo* passaging indicated that *S. pneumoniae* was recalcitrant to developing fluroquinolone resistance *in vivo*, we next sought to experimentally evolve the parental strain *in vitro* with increasing concentrations of either ciprofloxacin or levofloxacin. After 30 passages, TIGR4 did not develop significant resistance to ciprofloxacin as it only became resistant to 8 µg/mL (**Supplemental Figure 1**) whereas levofloxacin resistance was not observed after an extensive period of *in vitro* passaging (**Supplemental Figure 1**). Three independent colonies isolated from the *in vitro* passaging population under ciprofloxacin pressure were sequenced. Whole genome sequencing data demonstrated that those three colonies encode double point mutations in the on-target genes *parC* and *gyrA*, S79Y and S81F, respectively. The *in vitro* passaging isolates under levofloxacin pressure were not sequenced as no shifts in MIC were observed. These data suggest that acquisition of fluoroquinolone resistance by *de novo* mutation is a rare occurrence both *in vivo* and *in vitro*.

### Horizontal Acquisition of Fluoroquinolone Resistance

Being a naturally competent bacterial pathogen, antibiotic resistance in *S. pneumoniae* is oftentimes acquired via inter and intraspecies recombination events ^50^. We next sought to ascertain if horizontal transfer of on-target mutations conferring fluroquinolone resistance could be a source for acquisition of fluoroquinolone resistance in place of *de novo* mutation. To determine the relative frequency of horizonal transfer, the parental wild-type TIGR4 was transformed with PCR fragments encoding point mutations in either on-target genes, S81F in *gyrA* or S79Y in *parC*, and transformation efficiency was measured following selection on either levofloxacin (**Figure 2A**) or ciprofloxacin (**Figure 2J**). These mutations were selected due to their high prevalence in fluoroquinolone-resistant clinical isolates ^51–55^. As shown in **Figure 2A** and **J**, transformation with single on-target point mutations with the wild-type TIGR4 did not readily occur with many transformations failing to yield resistant colonies. When transformants were recovered, the transformation efficiency was significantly reduced compared to our positive control Tn-seq library whose efficiency was consistently an order of magnitude greater. Negative controls were assayed in parallel to account for spontaneous resistance mutations (**Figure 2A** and **J**). To determine whether this low transformation efficiency was strain or serotype specific, we also transformed additional *S. pneumoniae* strains including the D39 serotype 2 (**Figure 2D** and **M**) and CDC001 serotype 9V (**Figure 2G** and **P**) and observed similar transformation efficiencies. These data indicate that horizontal transfer for fluoroquinolone resistance as single on-target point mutations alone was relatively inefficient for acquisition of fluoroquinolone resistance compared to acquisition of spectinomycin resistance (Tn-Seq library).

**Figure 2.**
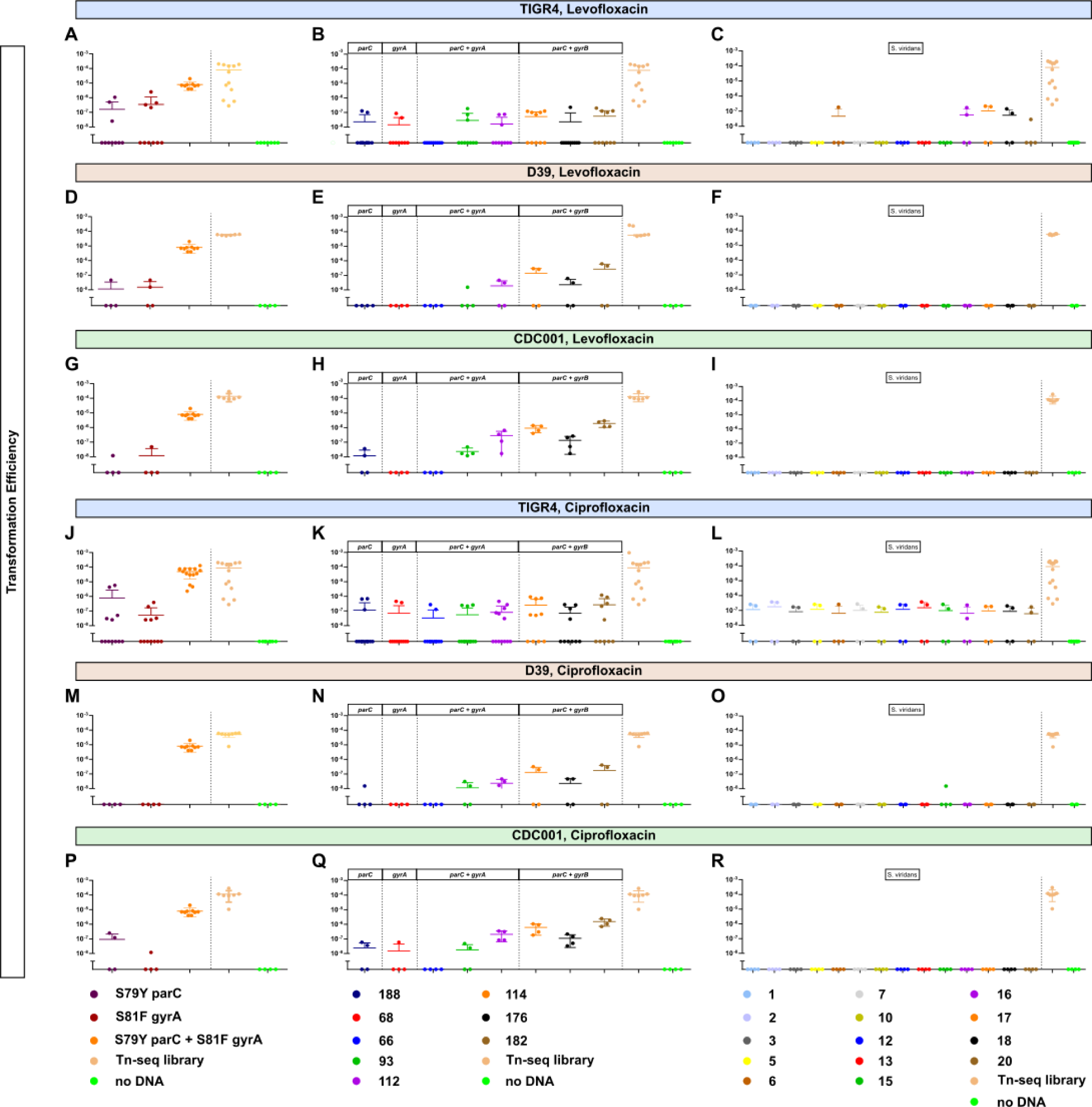
Transformation efficiency for fluoroquinolone resistance determinants across various strains. Transformation efficiency of fluroquinolone resistance determinants across multiple strain backgrounds including TIGR4, D39, and CDC001 (serotypes 4, 2, and 9V, respectively). DNA used for transformation included either PCR fragments encoding the respective mutations, (A, D, G, J, M, P), genomic DNA of *S. pneumoniae* clinical isolates harboring the respective mutations (B, E, H, K, N, Q), or genomic DNA of *S. viridans* clinical isolates harboring fluroquinolone resistance (C, F, I, L, O, R). Resistant colonies were recovered on 1X MIC of plates supplemented with either ciprofloxacin or levofloxacin. For all transformations, a Tn-seq library served as a positive control for competence. Transformation efficiency was calculated as the ratio of the number of transformants (CFUs/mL) selected on either levofloxacin or ciprofloxacin to the number of total bacteria (CFUs/mL). Each datapoint represents an individual biological replicate.

Compensatory mutations in the genome have been shown to ameliorate fitness costs imparted by on-target mutations, thereby facilitating the acquisition as well as allowing them to fixate in the population ^56, 57^. As such, conference of fluoroquinolone resistance, represented by enhanced transformation efficiency, may occur more readily when the donor DNA is from strains harboring fluoroquinolone resistance, which may contain compensatory mutations along with the on-target mutation. The transformation efficiency of TIGR4, D39, and CDC001 remained extremely low (**Figure 2B, E, H, K, N,** and **Q**). Transformation with genomic DNA of the resistant pneumococcal isolates that encoded double point mutations in both *parC* and *gyrA* occurred more often than those with genomic DNA that encoded only single point mutation in either *parC* or *gyrA*. Another major reservoir of antibiotic resistance determinants for *S. pneumoniae* is the viridans group streptococci ^58^. We next determined proclivity of *S. pneumoniae* to gain fluroquinolone resistance upon transformation with the interspecies genomic DNA from resistant *S. viridans* group clinical isolates, most of which encode double point mutations in both *gyrA* and *parC* except isolate #5 having triple point mutations in *gyrA*, *gyrB*, and *parC* (**Supplementary Table 3**). Transformation with genomic DNA of fluoroquinolone-resistant viridans group streptococci still rarely produced resistant strains. The TIGR4 strain had a relatively high efficiency of 10^-7^ when transformed with isolates 16, 17, and 18, which encoded double point mutations in *parC* and *gyrA,* under levofloxacin (**Figure 2C**) or when transformed with all isolates under ciprofloxacin (**Figure 2L**). No transformants were observed in the D39 and CDC001 strains (**Figure 2F, I, O,** and **R**). Taken together, infrequent transformation efficiencies with both intra- and inter-species genomic DNA encoding on-target mutations for fluoroquinolone resistance further corroborate acquisition of fluoroquinolone resistance determinants via horizontal transfer in *S. pneumoniae* is relatively challenging to achieve.

### Fitness Consequence of Fluoroquinolone Resistance Mutations

A key factor that determines whether a resistant strain will survive and disseminate in the environment or in the host is the fitness cost associated with the mutation(s) ^57^. Without antibiotic pressure, mutations that confer antibiotic resistance but carry high fitness costs are less likely to persist in the population ^37, 59^. In *S. pneumoniae*, fitness refers to the ability of the bacteria to colonize, replicate, invade different bodily organs, and disseminate to other hosts ^60^. We hypothesized that the absence of on-target mutations during the *in vivo* passaging experiment was potentially due to attenuation from the first step mutations conferring fluroquinolone resistance. To test this hypothesis, we generated single- and double-point mutants via transformation with a PCR fragment encoding the desired mutation(s). We successfully generated single point mutants S79Y *parC* and S81F *gyrA*, and double point mutants S79Y *parC* & S81F *gyrA* and S79Y *parC* & D435N *gyrB* via transformation with PCR fragments encoding specific mutations and confirmed the presence of those mutations without any other on-target mutations via whole genome sequencing and MIC testing (**Supplementary Table 2**). These mutations have been shown previously to confer high-level resistance to fluoroquinolones in clinical isolates ^61–64^. The growth rates of the mutants indicated that the single on-target mutants S79Y *parC* and S81F *gyrA* demonstrated a slight, but marked delay in growth compared to the wild-type TIGR4 (**Figure 3A**). Interestingly, the double mutant S79Y *parC* & S81F *gyrA* had similar growth rate to that of the wild-type, while the double mutant S79Y *parC* & D435N *gyrB* had the similar growth pattern to that of the S79Y *parC* mutant (**Figure 3A**).

**Figure 3.**
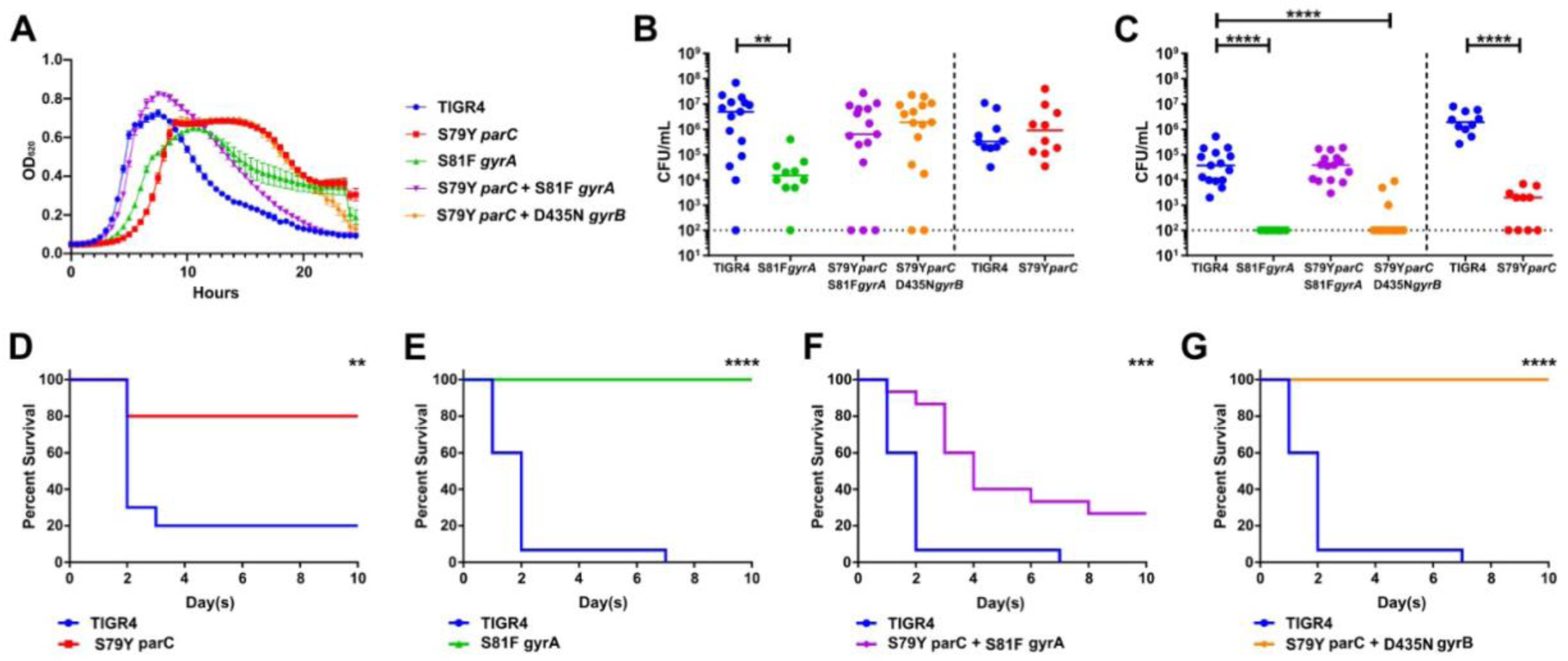
Individual fluroquinolone resistance mutations confer high fitness tradeoffs during invasive infection but not colonization. Respective mutants conferring fluroquinolone resistance were initially assayed for growth *in vitro* in semi-chemically defined media (A). Mice were infected intranasally with the parental TIGR4 wild-type or the isogenic fluroquinolone mutants. Nasal lavage at 24 hours (B) post-challenge was used to ascertain relative bacterial colonization burden. Invasive potential was assayed via blood titers at 24 hours (C) post- challenge. Infected mice were followed for ten days for relative survival between the parental and respective point mutants (D, E, F, G). The bacterial burden data were compared to that of the wild-type strain via Mann-Whitney and survival data were analyzed with Mantel-Cox log rank tests in Prism 7. *p < 0.05, **p < 0.01, ****p < 0.001. Bars represent mean, with each data point representing an individual mouse.

We next determined the *in vivo* fitness of the single and double mutants by determining their virulence in a murine model. For our pneumococcal virulence model, we intranasally infected mice with either the susceptible, parental TIGR4 strain or the various fluoroquinolone resistant mutants, followed by bacterial burden enumeration in the nasal lavage and blood as metrics of colonization and invasive capacity, respectively. Compared to the wild-type TIGR4, the single mutants in either *gyrA* or *parC* resulted in severe fitness defects *in vivo* (**Figure 3B- G**). Unlike the single mutant in *gyrA*, the single mutant in *parC* was able to colonize the nasopharynx as effectively as the wild-type (**Figure 3B**). Although there was a log decrease in bacterial burden compared to the wild-type, the double point mutant S79Y *parC* & S81F *gyrA* still colonized the nasopharynx (**Figure 3B**). Neither the single mutants S79Y *parC* or S81F *gyrA* were detected in the bloodstream, indicating that they were not able to disseminate and/or survive in the bloodstream to cause invasive disease (**Figure 3C**). When both the *gyrA* and *parC* mutations were present, fitness was restored to the equivalent to that observed in the parental strain in the blood, whereas the *parC & gyrB* double mutant retained attenuation similar to that observed in the single mutant (**Figure 3C**). In addition to measuring bacterial burden in the nasopharynx and blood, we monitored infected mice for survival for 10 days (**Figure 3D-G**). Mice infected with the single point mutants S79Y *parC* and S81F *gyrA* and the double point mutant S79Y *parC* & D435N *gyrB* showed a significant increase in survival compared to those infected with TIGR4 (**Figure 3D,E,** and **G**). Survival of mice infected with the single point mutants and the double points mutant S79Y *parC* & D435N *gyrB* was consistent with their undetectable bacterial burden in the blood. Mice infected with the double point mutant S79Y *parC* & S81F *gyrA* displayed significantly delayed mortality but overall survival was reduced, almost to the levels of mice infected with the parental strain (**Figure 3F**). These data suggest that the individual mutations associated with fluroquinolone resistance in either *gyrA* or *parC* impart significant fitness tradeoffs during infection that limited their emergence during our *in vivo* evolution experiment, demonstrating an important barrier to the acquisition of fluoroquinolone resistance in *S. pneumoniae*.

### Levofloxacin and host pressure favor emergence of tolerance over resistance

The defects in fitness observed in strains with on-target mutations conferring fluoroquinolone resistance could explain the absence of these on-target mutations and fluoroquinolone resistance in the experimentally evolved isolates. This led us to examine whether the isolates displayed tolerance phenotype as increasingly high dosages of levofloxacin failed to clear the infections. Tolerant strains, by definition, can survive transient exposure to high antibiotic concentrations that are otherwise lethal, thereby having slower kill kinetics than susceptible strains ^30^. We assayed the antibiotic kill kinetics of three experimentally evolved lineages and observed reduced antibiotic-mediated killing following addition of levofloxacin (**Figure 4A**), suggesting that tolerance to levofloxacin emerged in all three independent lineages. We next tested whether the experimentally evolved isolates conferred cross-tolerance to other clinically relevant fluoroquinolones, such as ciprofloxacin and moxifloxacin. Compared to the wild-type TIGR4, the isolates exhibited similar kill kinetics in response to both ciprofloxacin and moxifloxacin (**Figure 4B** and **C**). These data indicate that the evolved tolerance phenotype was specific for levofloxacin and not to other fluroquinolones.

**Figure 4.**
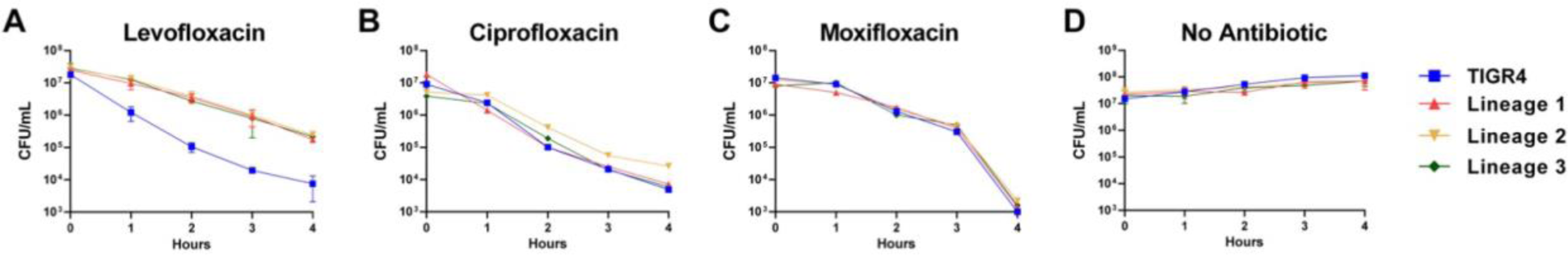
Experimentally evolved isolates of *S. pneumoniae* exhibited tolerant phenotype to levofloxacin. The TIGR4 wild-type and the final populations from the three independent lineages evolved in mice were assayed for time-kill kinetics in response to different fluroquinolone antibiotics- levofloxacin (A), ciprofloxacin (B), moxifloxacin (C), and no antibiotic (D). Data represents three biological replicates with the mean and standard deviation plotted.

Given that tolerance, rather than traditional resistance, emerged in the experimentally evolved isolates, we next undertook population analysis of any mutations that arose during *in vivo* passaging (**Figure 1**) to identify a possible genetic basis for the observed tolerance. From the whole genome sequencing data of the passaged isolates, we identified mutations as potential candidates for conferring tolerance based on the criteria that mutations arose in more than one independent lineage, were absent prior to *in vivo* passaging, and occurred in the bacterial population at high frequencies. Mutations in isochorismatase (A44D, A4D, and H172D) arose in three independent lineages during *in vivo* passaging with levofloxacin, starting in passage 8-10 and remaining in the population (**Figure 5**). In all lineages, loss of function mutations in *lctO* were identified; this gene encodes a lactate oxidase responsible for the conversion of pyruvate to lactate, generating hydrogen peroxide. Later passages revealed mutations in *spxB* (M1T, Q456*) and chlorohydrolase (Q17*, F19V) emerged and were sustained in the population until the final passage in two lineages (**Figure 5)**. The missense mutation in the start codon M1T of *spxB* results in a non-functional pyruvate oxidase, which, like lactate oxidase, is involved in pyruvate metabolism and hydrogen peroxide production. These data suggest that alterations in metabolic genes associated with pyruvate metabolism and generation of hydrogen peroxide may be one avenue for evolving tolerance to levofloxacin. We next assessed H_2_O_2_ production among the evolved isolates along with the parental TIGR4 and the Δ*spxB*Δ*lctO* double knockout, serving as positive and negative controls, respectively^49^.

**Figure 5.**
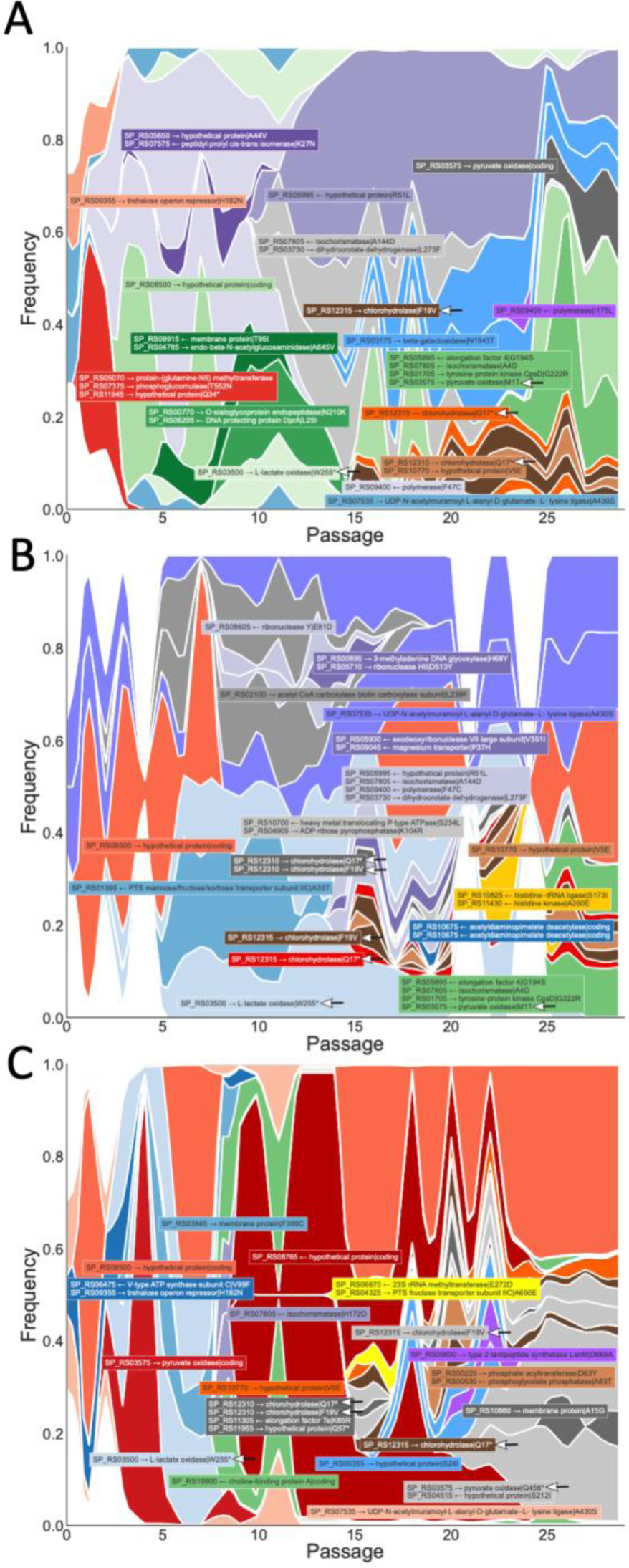
Population analysis of mutations arising during *in vivo* experimental evolution. Muller plots representing relative abundance of selected mutations that arose in multiple lineages during *in vivo* passaging with levofloxacin: Lineage 1 (A), Lineage 2 (B), Lineage 3 (C).

Compared to TIGR4, all three evolved isolates from the *in vivo* passaging produced significantly less H_2_O_2_ (**Supplemental Figure 2**). The loss of hydrogen peroxide production in the experimental evolved isolates suggests a prominent role for disruption of hydrogen peroxide production in tolerance to levofloxacin.

### Redox Stress Plays a Critical Role in Tolerance to Fluroquinolones

In the presence of fluoroquinolones, it is possible that the pneumococcus would be under redox stress from two major sources: their own H_2_O_2_ production and fluoroquinolone-induced radicals, as fluoroquinolones have been shown to exert redox stress, including the generation of hydroxyl radicals, on bacteria ^65^. To determine whether metabolic pathways responsible for the endogenously produced H_2_O_2_ contribute to a tolerance phenotype, we utilized the Δ*spxB*Δ*lctO* double knockout strain which abrogates pneumococcal H_2_O_2_ production (**Supplemental Figure 2**). The double mutant demonstrated reduced kill kinetics for both levofloxacin and ciprofloxacin (**Figure 6A** and **B**). As the production of H_2_O_2_ requires the presence of oxygen, we also undertook parallel time-kill experiments anaerobically as a control. Addition of either fluroquinolone under anaerobic condition resulted in no differences in kill kinetics between the parental wild-type and the Δ*spxB*Δ*lctO* double knockout strain (**Figure 6C** and **D**). These data indicate that tolerance is linked to the metabolic capacity of the pneumococcus to generate hydrogen peroxide.

**Figure 6.**
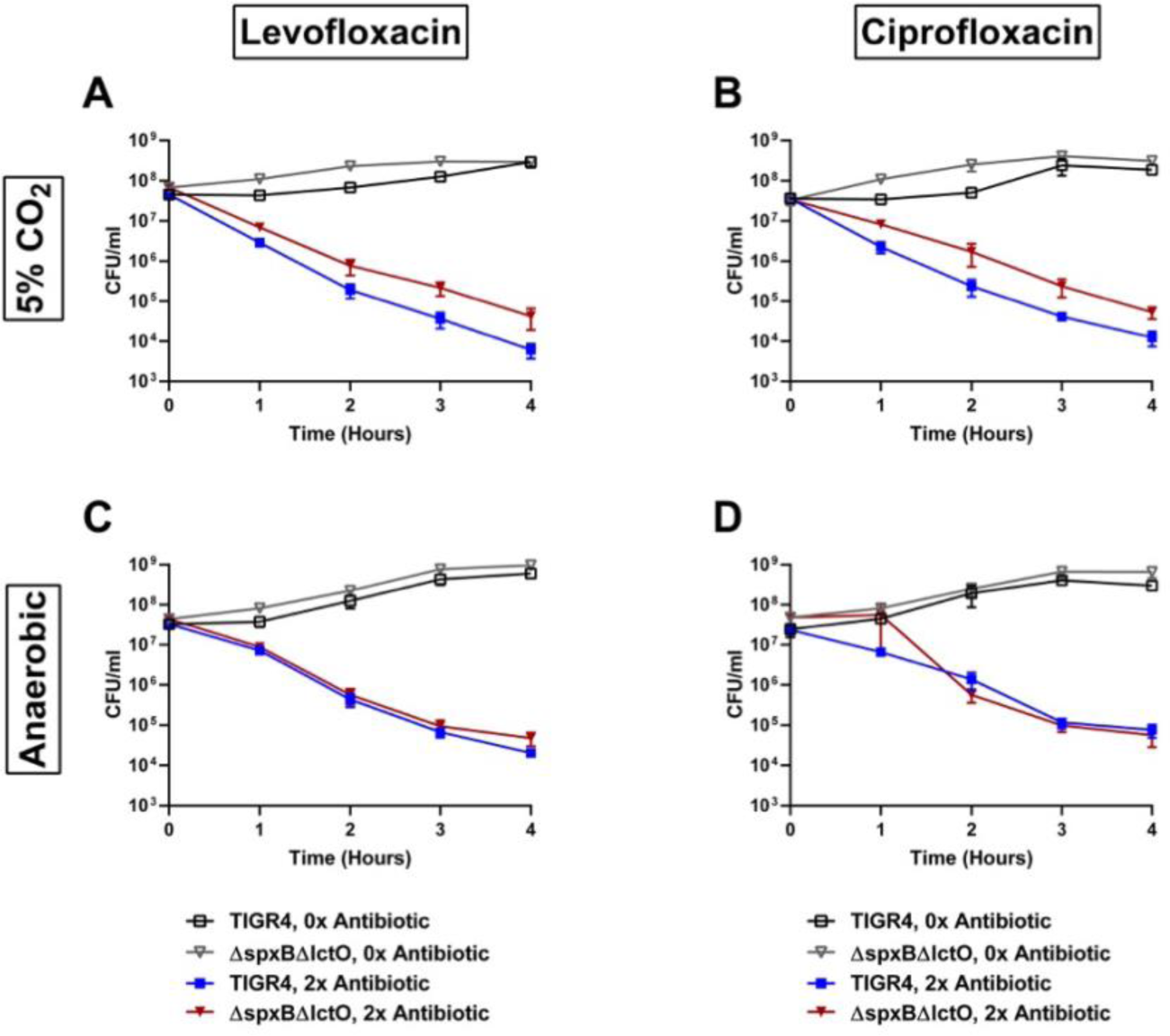
Disruption of endogenous hydrogen peroxide production confers tolerance to fluroquinolones. The TIGR4 wild-type and the Δ*spxB*Δ*lctO* double knockout strain were assayed for time-kill kinetics in response to different fluroquinolone antibiotics and levels of oxygen: levofloxacin (A), ciprofloxacin (B), levofloxacin under anaerobic conditions (C), and ciprofloxacin under anaerobic conditions (D). Data represents three biological replicates with the mean and standard deviation plotted.

Besides the H_2_O_2_ endogenously produced by the pneumococcus, the bactericidal activities of fluroquinolones may be impacted by the free radicals induced by the antibiotic. To test this, we utilized a clinically approved drug (edaravone) that scavenges free radicals to determine the effect of diminished free radicals in the antibiotic kill kinetic experiments ^66, 67^.

Supplementation of the media with edaravone dramatically reduced the bactericidal activities of both levofloxacin and ciprofloxacin, resulting in over two orders of magnitude reduction in killing following four hours of antibiotic exposure (**Figure 7A** and **B**). This result suggests that, in the absence of fluoroquinolone-induced free radicals, the pneumococcus can become tolerant to fluoroquinolone killing. As both redox stress and fluroquinolones can result in DNA damage and fragmentation that can contribute to bactericidal activity ^68, 69^, we next sought to determine whether the protection engendered by edaravone correlated with a corresponding decrease in DNA fragmentation during fluroquinolone treatment. While treatment with either levofloxacin or ciprofloxacin resulted in elevated DNA fragmentation (**Figure 7C** and **D**), this effect was completely abrogated upon treatment with edaravone, bringing the antibiotic exposed cells to basal wild-type levels (**Figure 7C** and **D**). Additional controls for the TUNEL assay are shown in **Supplemental Figure 3**. These data further substantiate a critical role of both endogenously produced or fluoroquinolone- induced hydroxyl radicals on the kinetics of fluroquinolone mediated bactericidal activity on the pneumococcus.

**Figure 7.**
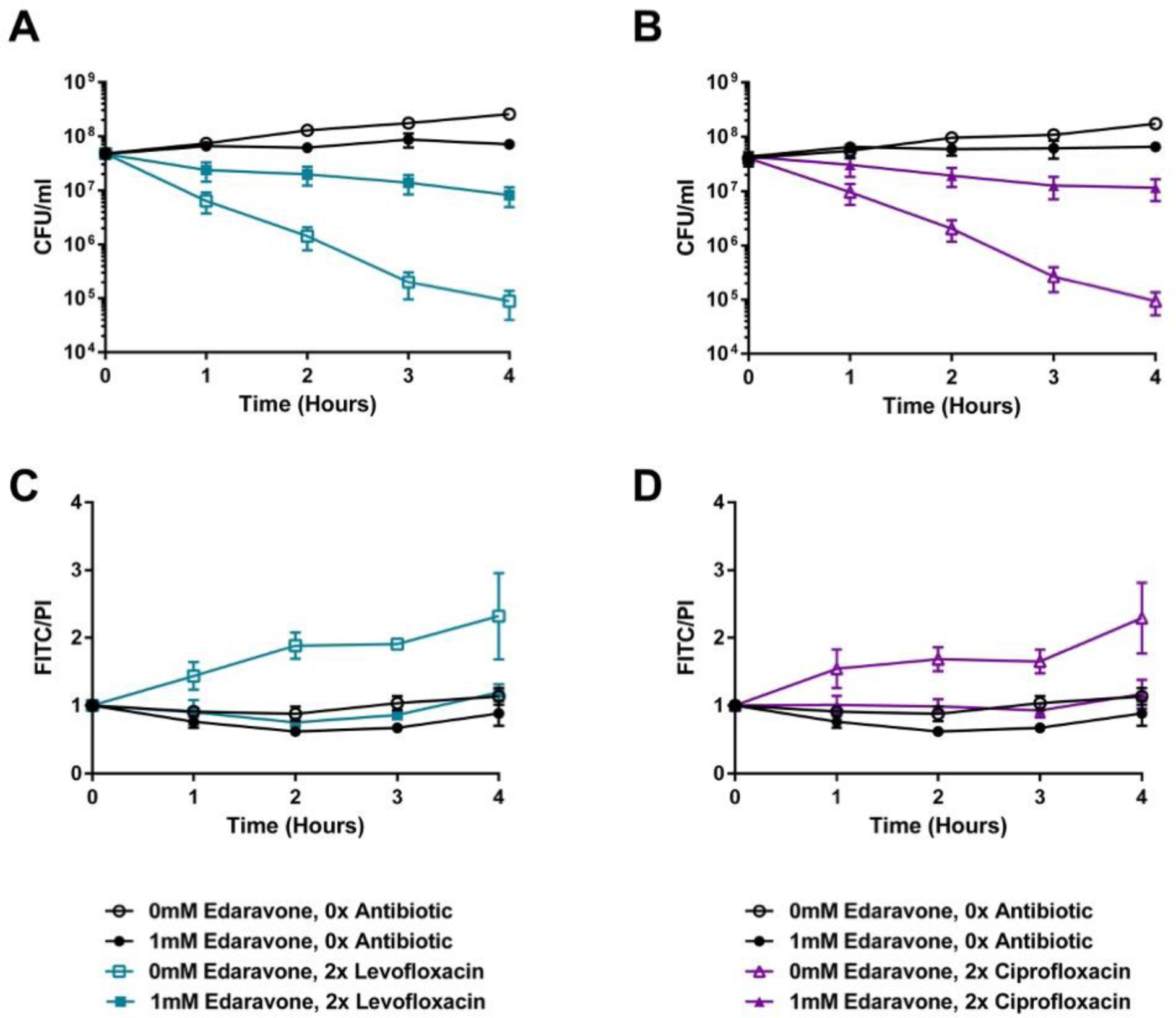
*S. pneumoniae* treated with free-radical scavenger edaravone demonstrates tolerance to fluoroquinolones and decreased DNA fragmentation. The TIGR4 wild-type was assayed for time-kill kinetics upon edaravone supplementation in the presence of levofloxacin (A) or ciprofloxacin (B). DNA fragmentation in response to either levofloxacin (C) or ciprofloxacin (D) was measured concurrently with the time-kill kinetic experiment. Data represents three biological replicates with the mean and standard deviation plotted.

### Tolerance Promotes Survival During Antibiotic Treatment of Infection

These data suggest that, during infection, mutations that reduce stress by oxygen radicals whilst retaining fitness may be one strategy whereby the pneumococci could evade the bactericidal activities of fluroquinolones through tolerance, leading to antibiotic treatment failure. To test this, a competitive index experiment was undertaken whereby mice were infected with a 1:1 ratio of wild-type and the Δ*spxB*Δ*lctO* double knockout strain. Bacterial burden for each individual mouse was followed prior to and following antibiotic treatment with levofloxacin (**Supplemental Figure 4**). When levofloxacin was not administered (**Figure 8A**), no differences in competitive index between the wild-type and double mutant were observed. Likewise, immediately prior to treatment, no significant differences in competitive index were noted (**Figure 8B**). Starting at 2 hours and continuing up to 10 hours post levofloxacin treatment, the Δ*spxB*Δ*lctO* double knockout strain demonstrated a significant fitness advantage over the parental wild-type strain (**Figure 8B**). These data suggest that metabolic solutions conferring antibiotic tolerance can lead to antibiotic treatment failure in the absence of traditional resistance mechanisms.

**Figure 8.**
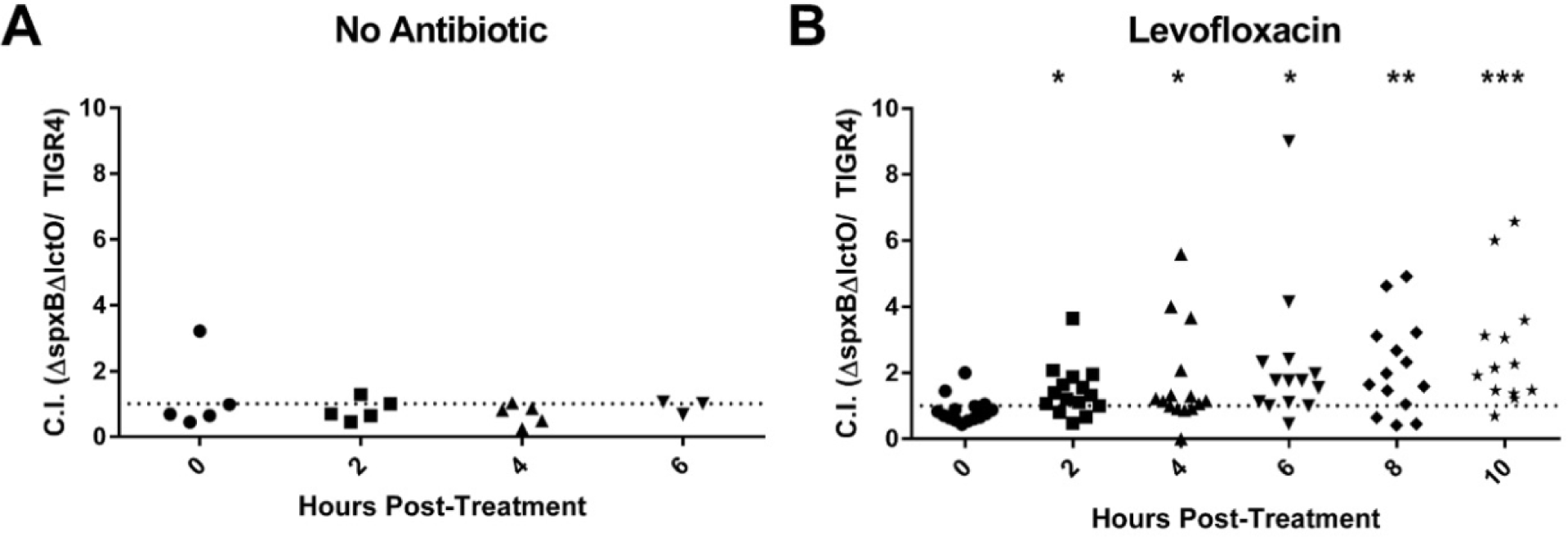
*S. pneumoniae* strains lacking hydrogen peroxide production outcompetes wild-type upon treatment with levofloxacin. Mice were infected with TIGR4 wild-type and the Δ*spxB*Δ*lctO* double knockout strain and competitive indexes of the Δ*spxB*Δ*lctO* double knockout strain without antibiotic treatment (A) and following treatment with levofloxacin (B) were calculated. Competitive index of timepoints 2 through 10 hours was compared to that of timepoint 0 for each condition using unpaired parametric t-test in Prism 7. *p < 0.05, **p < 0.01, ***p < 0.001. Each data point represents an individual mouse that was tracked consecutively over the respective time points.

## Discussion

In this study, we aimed to investigate the discrepancy between high usage rate of fluoroquinolones and the low resistance rates observed in *S. pneumoniae*. We found that the single point mutations in either of the on-target enzymes, topoisomerase IV *parC* and DNA gyrase *gyrA*, conferred severe fitness costs for invasive disease unless a second point mutation was acquired simultaneously. This agrees with clinical studies report that high-level fluoroquinolone resistant *S. pneumoniae* isolates often harbor double point mutations in *parC* and *gyrA* ^70–72^. The presence of the on-target double point mutations in clinical settings is consistent with our finding that the double point mutants exhibited high resistance potential with low fitness costs, while maintaining a virulence level similar to that of the wild-type ^73^. Rozen and colleagues also reported that double mutations in *parC* and *gyrA* not only increased resistance potential, but also lessened the *in vitro* fitness costs associated with fluoroquinolone resistance mutations ^74^. Our results do not preclude the presence of additional compensatory mechanisms, particularly for invasive clinical isolates for which only single mutations in either *parC* or *gyrA* have been identified ^61, 75–77^. The high fitness cost imparted by the individual mutations conferring fluoroquinolone resistance may partly explain the low prevalence of fluoroquinolone resistance, as both mutations would need to be acquired simultaneously to avoid deleterious fitness tradeoffs.

In agreement with the steep fitness costs associated with *de novo* generation of fluroquinolone resistance, repeated exposure during *in vivo* infection did not result in the emergence of mutations in either *gyrA* or *parC*. This agrees with previous studies in rabbit models whereby repeated exposures to either levofloxacin or moxifloxacin did not result in appreciable shifts in *S. pneumoniae* recovered from the infection site ^78^. Despite the lack of appreciable shifts in MIC, the experimentally evolved strains resulted in phenotypic tolerance to levofloxacin, whereby markedly reduced bactericidal kill kinetics were observed. This underscores the importance of such off-target mutations that modulate antibiotic kill kinetics as a potential driver of antibiotic treatment failure. As antibiotics can induce profound metabolic disruptions contributing to their bactericidal activity ^79^, mutations of metabolic gene networks to facilitate tolerance to antibiotics is a common mechanism adopted by multiple bacterial pathogens in response to different classes of antibiotics. Perturbation of fatty acid metabolic networks have been demonstrated to induce an antibiotic tolerant state in pathogenic Mycobacterial species, suggesting important roles for carbon source utilization in antibiotic efficacy ^80^. Zhang and colleagues demonstrated that activated PpnN (YgdH) in *E. coli* participated in the inhibition of nucleotides synthesis and increase in degradation, conferring tolerance to ciprofloxacin and ofloxacin ^81^. PpnN is a nucleosidase that plays a role in the metabolism of purine nucleotides; therefore, it is not a direct target of fluoroquinolones. Not only does PpnN confer tolerance to fluoroquinolones, but it also offers fitness advantages during stringent responses to stresses ^81^. Mutations that enhance cytoplasmic acidification and subsequent shutdown of protein synthesis can promote the emergence of persister cell states^82^. These data suggest that alteration of metabolic networks is an important aspect of the capacity of the pathogen to evade antibiotic mediated killing.

In our study, the population sequencing and functional characterization of the evolved populations indicated an important role of hydroxyl radicals for the observed tolerance phenotypes, a finding subsequently confirmed via targeted mutagenesis and chemical inhibition. This is in strong agreement with previous observations, whereby reactive oxygen species production is a critical aspect of the post-antibiotic effect for delayed regrowth of fluroquinolone treated *S. pneumoniae* ^69^. Additional support for the role of oxidative stress in potentiation of fluroquinolone killing comes from the observation that under levofloxacin stress, the pneumococcus upregulates genes involved in iron transport, whose substrate can subsequently generate oxidative stress through the Fenton reaction ^83^. Ciprofloxacin, along with penicillin and kanamycin, have been demonstrated to increase the intracellular reactive oxygen species detected in *S. pneumoniae* ^84^. Reduction of oxidative stress has been shown to promote bacterial survival upon exposure to bactericidal antibiotics ^85^. Moreover, the observation that scavenging intracellular oxidative molecules via edaravone treatment confers a tolerance phenotype while reversing the DNA fragmentation provides additional evidence that DNA fragmentation is an important aspect of the bactericidal activities of fluroquinolones ^85^. These data also suggest that unforeseen activities of non-antibiotic drugs may have unintended consequences on bacterial physiology that reduce antibiotic efficacy. Despite the importance of redox active pathways for the activity of fluoroquinolone mediated killing, robust bactericidal activity still was observed under anaerobic conditions, indicating that distinct mechanisms dependent on the specific growth conditions are likely involved. These data shown here further support the importance of redox stress for the bactericidal activity of fluroquinolones and suggest that mutations in metabolic networks represent one pathway by which pathogens can evade antibiotic therapy.

Since the metabolic adaptations that facilitate antibiotic tolerance may lead to antibiotic treatment failure and facilitate the acquisition of high-level resistance, targeting such pathways may prove useful avenues for more effective treatment strategies. There is increasing recognition of the collateral consequences of antibiotic resistance acquisitions being a viable target for combination antibiotic therapy. Detailed analysis of the fitness costs of antibiotic resistance can provide insight into unique genetic and phenotypical liabilities that can be therapeutically targeted ^86^. Identification of highly conserved pathways that can be therapeutically targeted to potentiate the activity of antibiotics, particularly against tolerant populations, provides avenues to expand the therapeutic lifespan of current antibiotics ^87^. Our findings suggest metabolic pathways that increase cellular redox stress may prove to be attractive candidates for such approaches against *S. pneumoniae* infections.

## Acknowledgements

JWR and work described herein was supported in part by 1U01AI124302 and by ALSAC.

## Supplementary Material

**Supplemental Table 1.** Prevalence of fluoroquinolone resistance from 1988 to 2015

| Fluoroquinolone Usage<br>(Prescriptions per 1000<br>Persons) | Fluoroquinolone Prevalence | Periods | Region | Ref |
| --- | --- | --- | --- | --- |
| 37.5 | 1.2% isolates first step mutation in *parC*<br>6.7% efflux phenotype | 2002-2006 | Germany | 1 |
| Not available | 0.2% resistant to levofloxacin | 2004-2005 to 2014-2015 | Germany | 2 |
| Ciprofloxacin<br>38.2 to 59.4 | 1.8% resistant to ciprofloxacin | 1988-2006 | Canada | 3-5 |
| Respiratory fluoroquinolone<br>5.3 to 27.4 | 1.2% non-susceptible to levofloxacin<br>1.1% non-susceptible to moxifloxacin | 1988-2006 | Canada | 3-5 |
| Not available | 0% in 1993 to 1.7% in 1997 and 1998 | 1988 to 1997 | Canada | 6 |
| Not available | 1.5%<br>2.9% | 1993 to 1994<br>1997 to 1998 | Canada | 1 |
| Not available | <2% resistant to levofloxacin | 2007-2016 | Canada | 4,7 |
| 39 to 106 | ≤ 2% resistant to levofloxacin<br>≤ 1% resistant to moxifloxacin | 1995-2002<br>2008 | USA | 8,9 |
| Not available | 4-13% resistant to fluoroquinolones (ciprofloxacin) | 2000 | Croatia and Hong Kong | 10 |
| Not available | 4% resistant to newer fluoroquinolones (levofloxacin, moxifloxacin) | 2009 to 2010 | Asia | 11 |
| Not available | 7.1% | 1998-1999 | Spain | 12 |
| Not available | 1.64 % resistant to ciprofloxacin, 0% to other fluoroquinolones | 2000-2002 | Hungary | 13 |

**Supplemental Table Error!.**
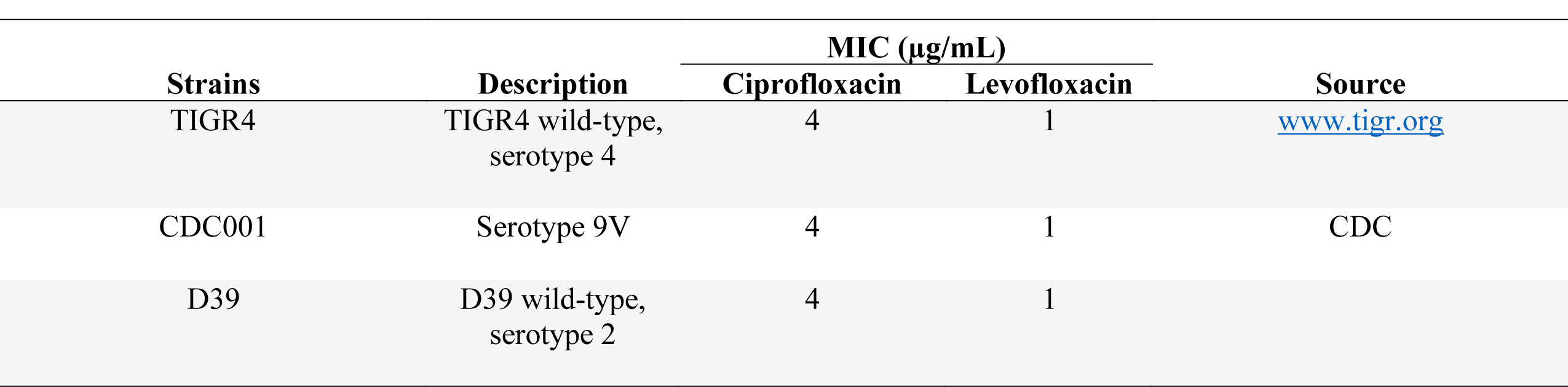
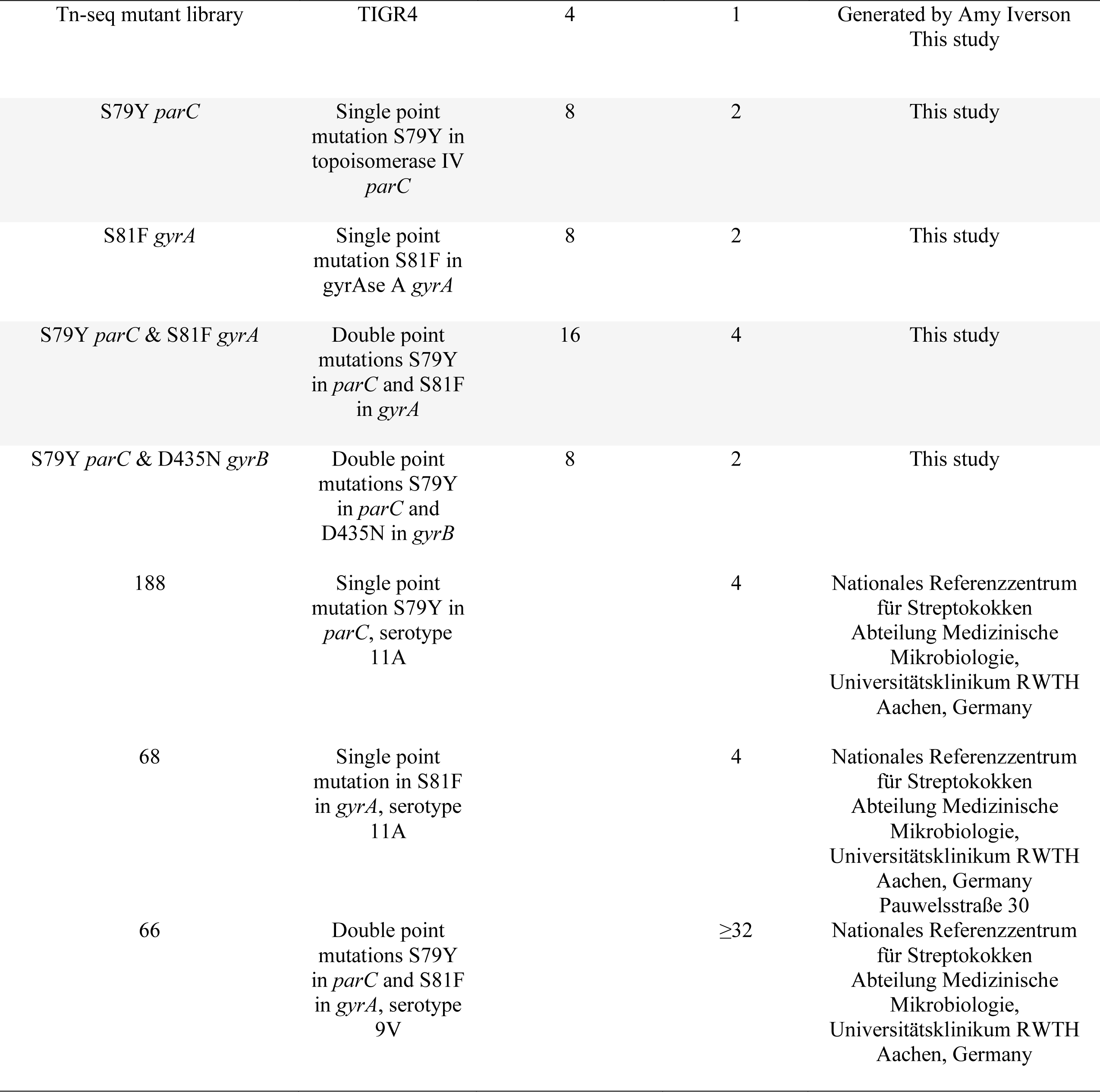
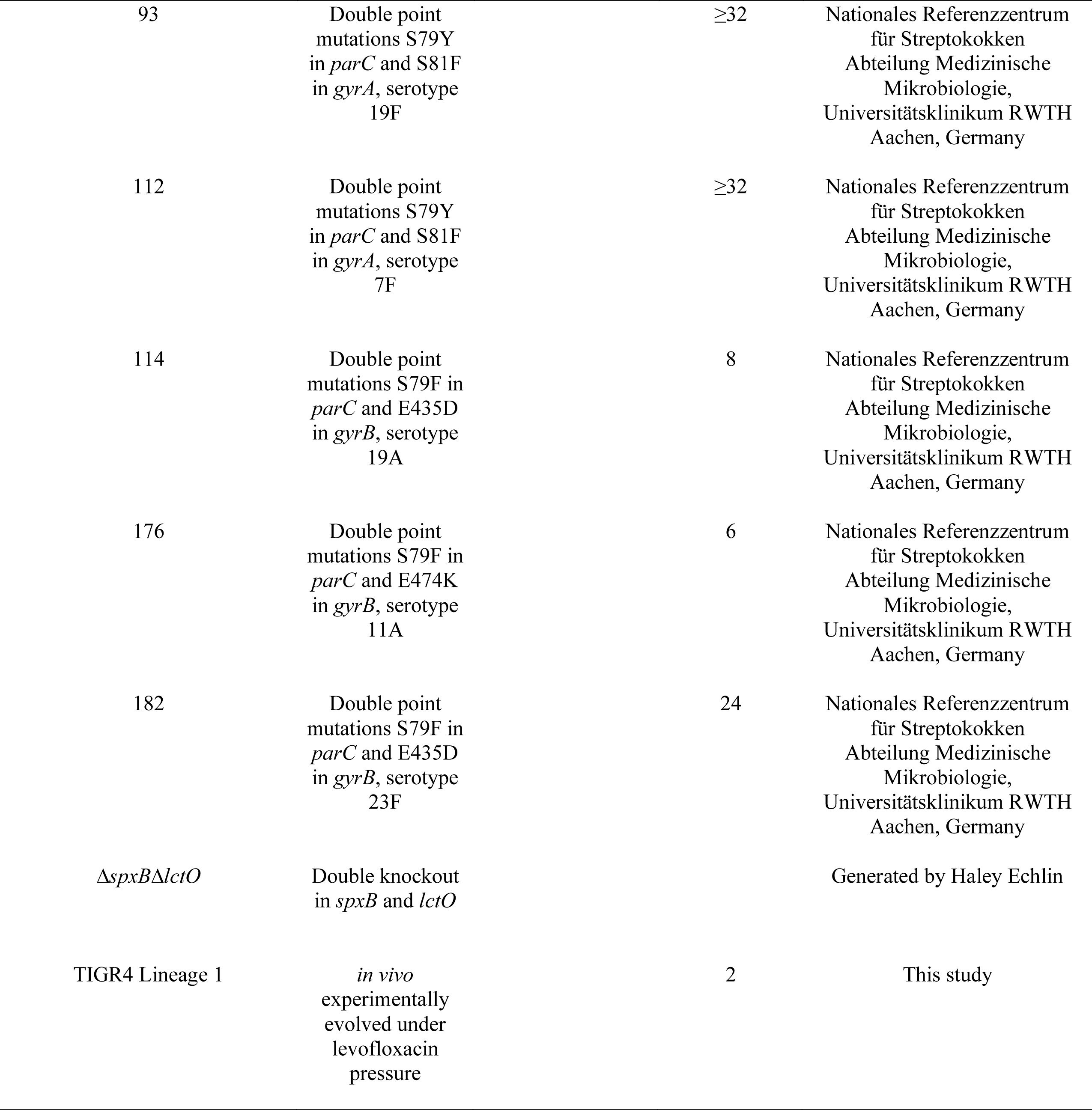
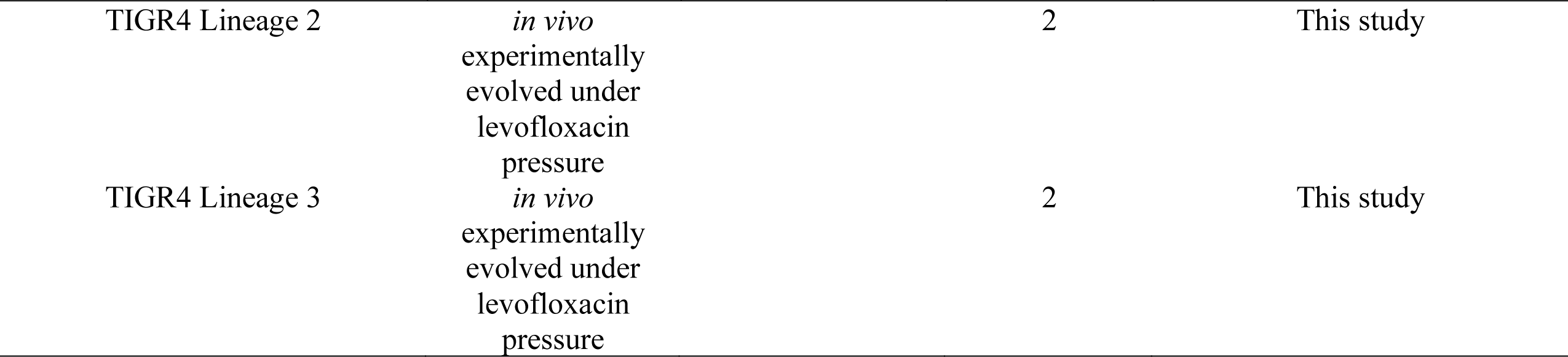
No text of specified style in document.. Strains used in this study and respective MICs

**Supplemental Table 3.** *S. viridans* strains used in this study and respective MICs

| Strain | Description | Levofloxacin<br>MIC (µg/mL) | Source |
| --- | --- | --- | --- |
| 1 | Double point mutations S79Y in *parC* and S81F in *gyrA* | 64 | SJCRH Clinical Microbiology<br>Laboratory |
| 2 | Double point mutations S79Y in *parC* and S81F in *gyrA* | 16 | SJCRH Clinical Microbiology<br>Laboratory |
| 3 | Double point mutations S79Y in *parC* and S81F in *gyrA* | 64 | SJCRH Clinical Microbiology<br>Laboratory |
| 5 | Triple mutations S79Y in *parC*, S81F in *gyrA*, and D474E<br>in *gyrB* | 16 | SJCRH Clinical Microbiology<br>Laboratory |
| 6 | Double point mutations S79Y in *parC* and S81F in *gyrA* | 16 | SJCRH Clinical Microbiology<br>Laboratory |
| 7 | Double point mutations S79Y in *parC* and S81F in *gyrA* | 16 | SJCRH Clinical Microbiology<br>Laboratory |
| 10 | Double point mutations S79Y in *parC* and S81F in *gyrA* | 64 | SJCRH Clinical Microbiology<br>Laboratory |
| 12 | Double point mutations S79Y in *parC* and S81F in *gyrA* | 16 | SJCRH Clinical Microbiology<br>Laboratory |
| 13 | Double point mutations S79Y in *parC* and S81F in *gyrA* | 32 | SJCRH Clinical Microbiology<br>Laboratory |
| 15 | Double point mutations S79Y in *parC* and S81F in *gyrA* | 16 | SJCRH Clinical Microbiology<br>Laboratory |
| 16 | Double point mutations S79Y in *parC* and S81F in *gyrA* | 64 | SJCRH Clinical Microbiology Laboratory |
| 17 | Double point mutations S79Y in *parC* and S81F in *gyrA* | 16 | SJCRH Clinical Microbiology Laboratory |
| 18 | Double point mutations S79Y in *parC* and S81F in *gyrA* | 16 | SJCRH Clinical Microbiology Laboratory |
| 20 | Double point mutations S79Y in *parC* and S81F in *gyrA* | 16 | SJCRH Clinical Microbiology Laboratory |

**Supplementary Figure 1.**
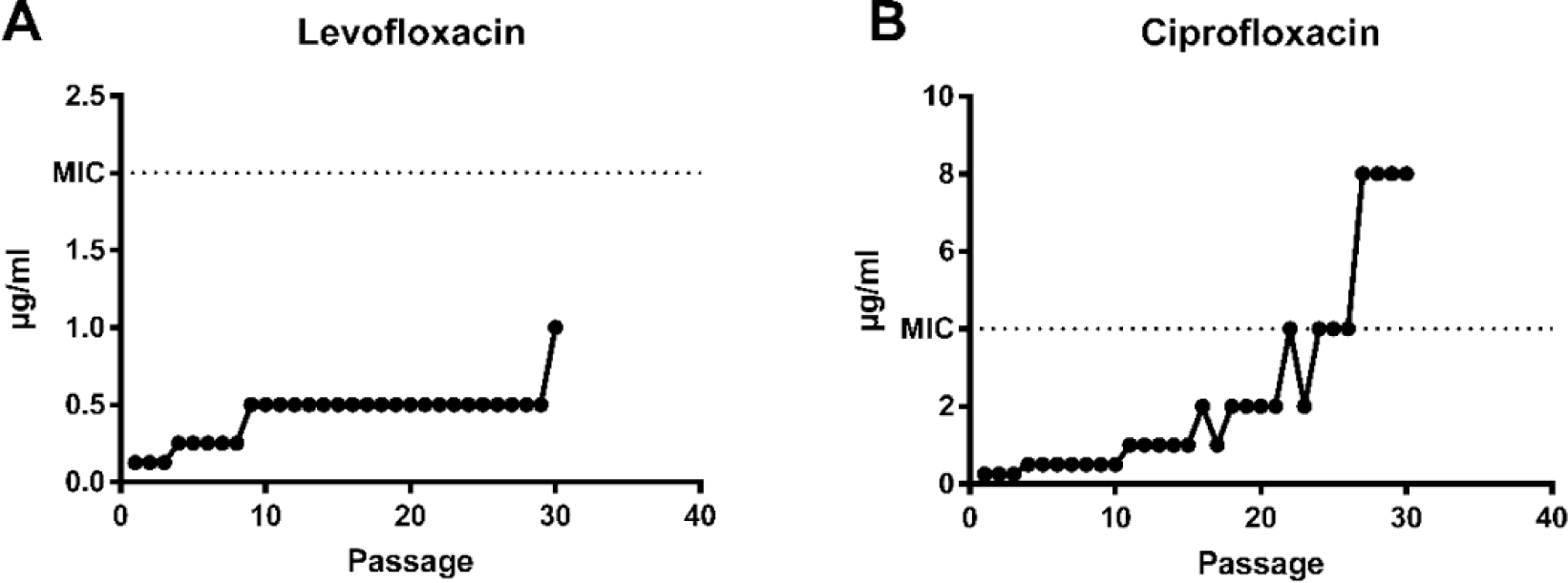
*S. pneumoniae* demonstrated *de novo* resistance *in vitro in vitro* passaging. *S. pneumoniae* was exposed to gradually increasing concentrations of both **A)** levofloxacin and **B)** ciprofloxacin for 30 days.

**Supplementary Figure 2.**
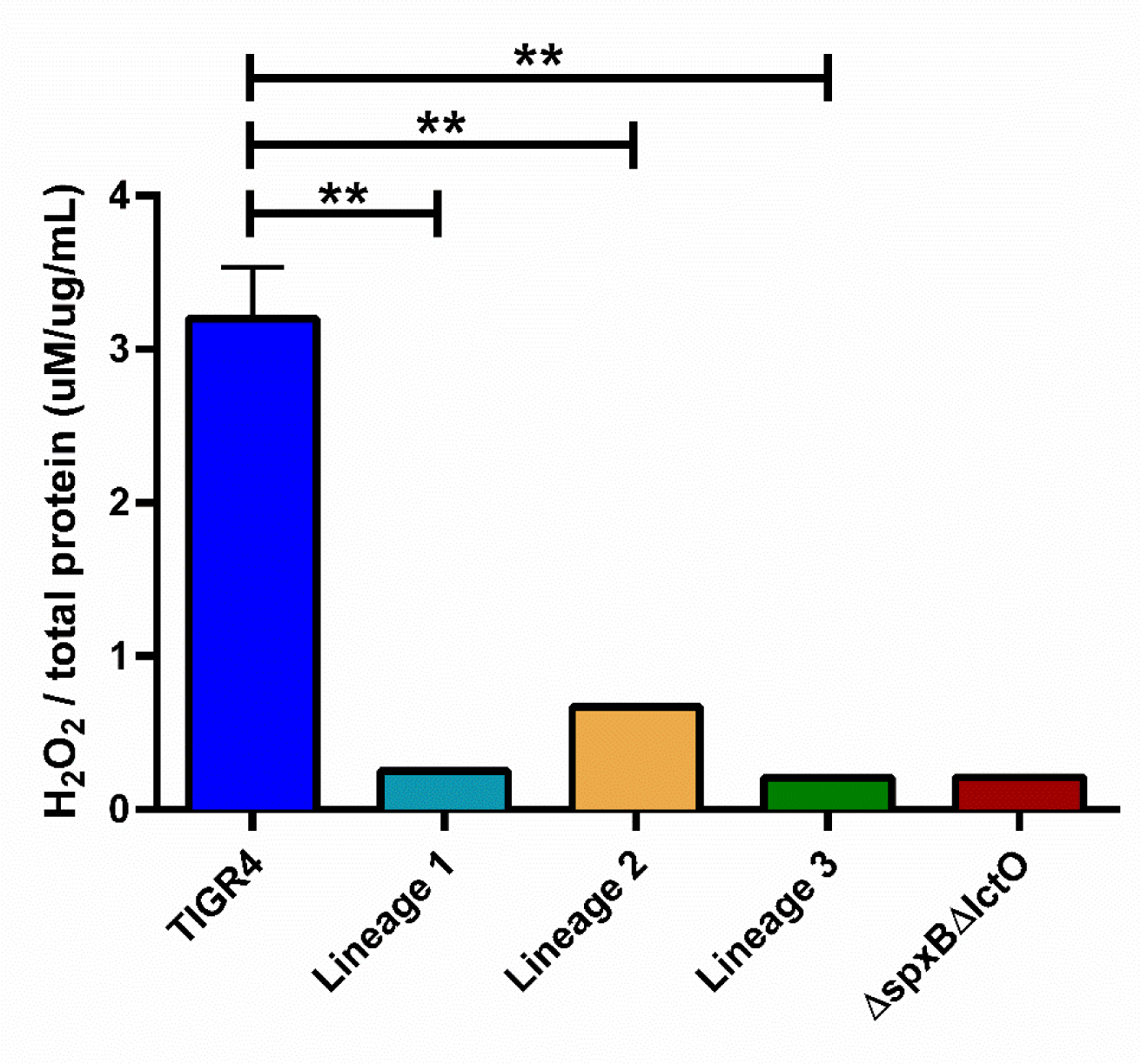
The evolved isolates produced minimal H_2_O_2_. H_2_O_2_ production was measured using the Amplex Red kit. Strains included the wild-type TIGR4, passage 29 of three independent lineages generated through *in vivo* passaging under levofloxacin treatment, and the Δ*spxB*Δ*lctO* double knockout mutant. The Δ*spxB*Δ*lctO* double knockout mutant served as the negative control. Amount of H_2_O_2_ was normalized to amount of protein each respective strain produced,determined via BCA assay. Hydrogen peroxide production of the evolved isolates was compared to that of wild-type TIGR4 via unpaired parametric t-test in Prism 7. **p-value ≤ 0.01.

**Supplemental Figure 3.**
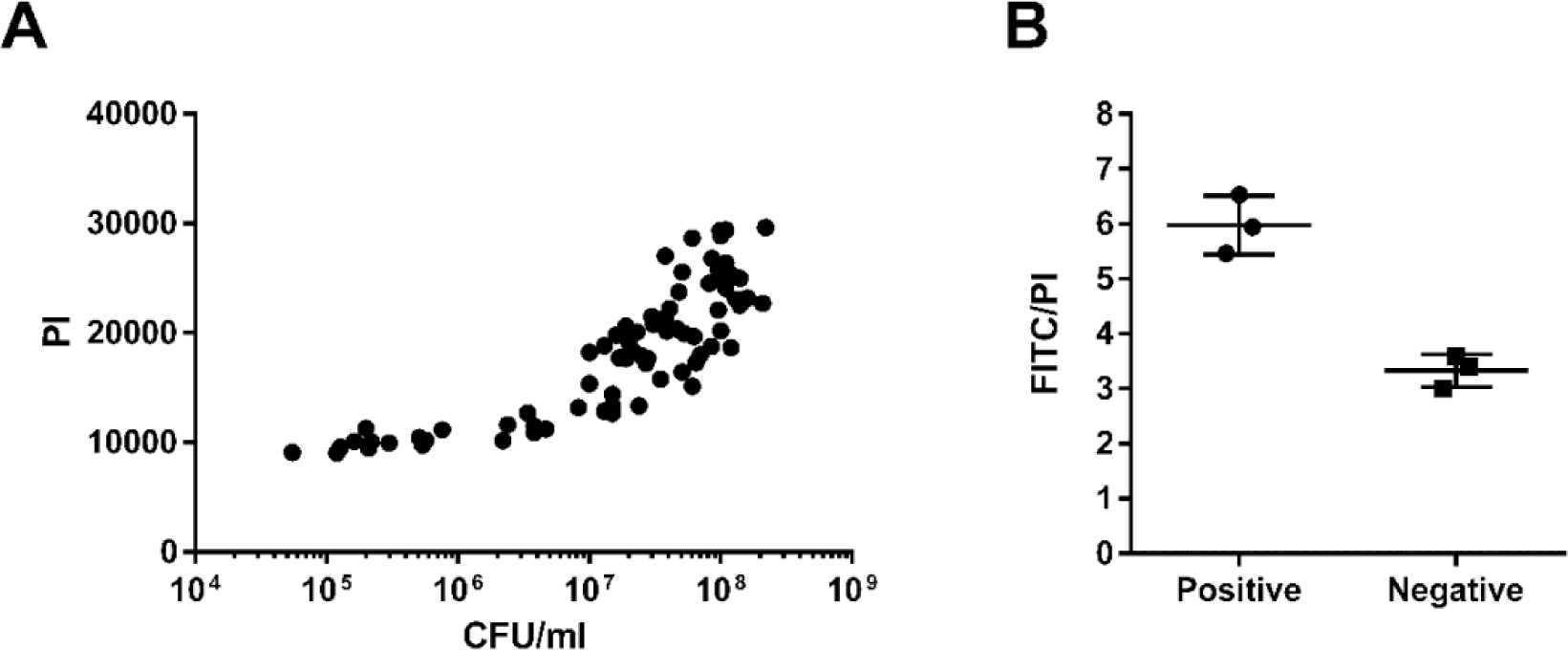
Controls for TUNEL assay. Correlation between PI values and CFU/mL of 5 mL cultures used for TUNEL staining (A). Ratio of FITC to PI levels of the positive and negative controls provided by the kit, detected at the same time as *S. pneumoniae* fixed cells (B).

**Supplemental Figure 4.**
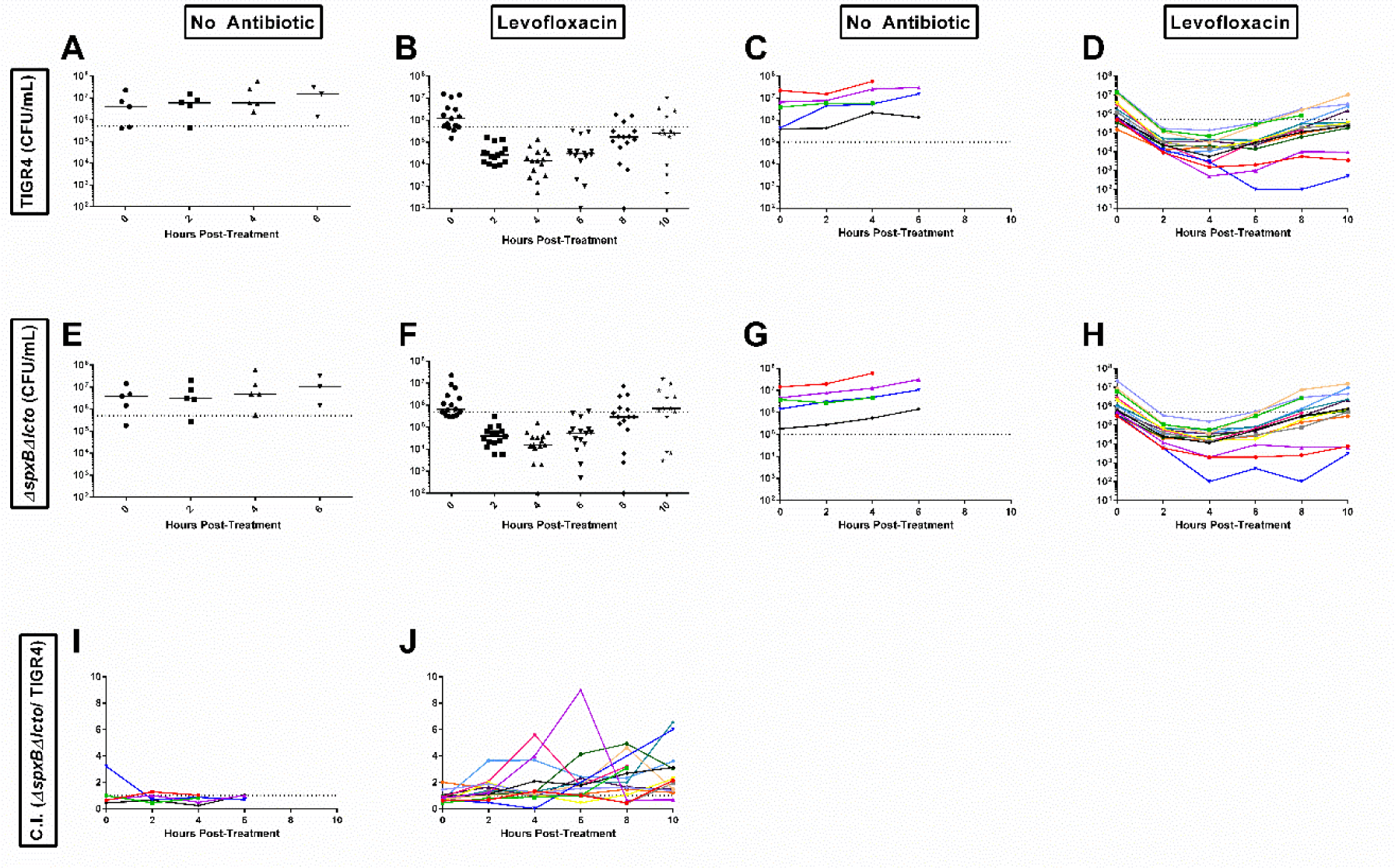
Total CFU/mL numbers and competitive index following each mouse. CFU/mL of TIGR4 was calculated as the CFU/mL on the plates containing 20 µg/mL neomycin minus the CFU/mL on the plates containing 20 µg/mL neomycin plus 1 µg/mL erythromycin for treatment with PBS (A) or levofloxacin (B). The bacterial burden in the blood of each mouse infected with TIGR4 was followed post-treatment with either PBS (C) or levofloxacin (D). CFU/mL of the Δ*spxB*Δ*lctO* double mutant was the CFU/mL on the plates containing 20 µg/mL neomycin plus 1 µg/mL erythromycin for treatment with PBS (E) or levofloxacin (F). The bacterial burden in the blood of each mouse infected with Δ*spxB*Δ*lctO* was followed post-treatment with either PBS (G) or levofloxacin (H). The competitive index of Δ*spxB*Δ*lctO* over TIGR4 of each infected mouse was followed post-treatment with either PBS (I) or levofloxacin (J).

